# Flow-cytometric microglial sorting coupled with quantitative proteomics identifies moesin as a highly-abundant microglial protein with relevance to Alzheimer’s disease

**DOI:** 10.1101/802694

**Authors:** Sruti Rayaprolu, Tianwen Gao, Hailian Xiao, Supriya Ramesha, Laura D. Weinstock, Jheel Shah, Duc M. Duong, Eric B. Dammer, James A. Webster, James J. Lah, Levi B. Wood, Ranjita Betarbet, Allan I. Levey, Nicholas T. Seyfried, Srikant Rangaraju

**Affiliations:** Department of Neurology, Emory University, Atlanta, GA 30322, USA; Department of Biochemistry, Emory University, Atlanta, GA 30322, USA; School of Medicine, Emory University, Atlanta, GA 30322, USA; Xiangya Hospital, Central South University, Changsha, Hunan, 410008, China; Parker H. Petit Institute for Bioengineering and Bioscience, Wallace H. Coulter Department of Biomedical Engineering, and Georgia W. Woodruff School of Mechanical Engineering, Georgia Institute of Technology, Atlanta, GA, 30332 USA

**Keywords:** microglia, proteomics, mass spectrometry, FACS, MACS, Alzheimer’s disease

## Abstract

**Background:** Proteomic characterization of microglia provides the most proximate assessment of functionally relevant molecular mechanisms of neuroinflammation. However, microglial proteomics studies have been limited by low cellular yield and contamination by non-microglial proteins using existing enrichment strategies.

**Methods:** We coupled magnetic-activated cell sorting (MACS) and fluorescence activated cell sorting (FACS) of microglia with tandem mass tag-mass spectrometry (TMT-MS) to obtain a highly-pure microglial proteome and identified a core set of highly-abundant microglial proteins in adult mouse brain. We interrogated existing human proteomic data for Alzheimer’s disease (AD) relevance of highly-abundant microglial proteins and performed immuno-histochemical and *in-vitro* validation studies.

**Results:** Quantitative multiplexed proteomics by TMT-MS of CD11b+ MACS-enriched (*N* = 5 mice) and FACS-isolated (*N* = 5 mice), from adult wild-type mice, identified 1,791 proteins. A total of 203 proteins were highly abundant in both datasets, representing a core-set of highly abundant microglial proteins. In addition, we found 953 differentially enriched proteins comparing MACS and FACS-based approaches, indicating significant differences between both strategies. The FACS-isolated microglia proteome was enriched with cytosolic, endoplasmic reticulum, and ribosomal proteins involved in protein metabolism and immune system functions, as well as an abundance of canonical microglial proteins. Conversely, the MACS-enriched microglia proteome was enriched with mitochondrial and synaptic proteins and higher abundance of neuronal, oligodendrocytic and astrocytic proteins. From the 203 consensus microglial proteins with high abundance in both datasets, we confirmed microglial expression of moesin (Msn) in wild-type and 5xFAD mouse brains as well as in human AD brains. Msn expression is nearly exclusively found in microglia that surround Aβ plaques in 5xFAD brains. In *in-vitro* primary microglial studies, Msn silencing by siRNA decreased Aβ phagocytosis and increased lipopolysaccharide-induced production of the pro-inflammatory cytokine, tumor necrosis factor (TNF). In network analysis of human brain proteomic data, Msn was a hub protein of an inflammatory co-expression module positively associated with AD neuropathological features and cognitive dysfunction.

**Conclusions:** Using FACS coupled with TMT-MS as the method of choice for microglial proteomics, we define a core set of highly-abundant adult microglial proteins. Among these, we validate Msn as highly-abundant in plaque-associated microglia with relevance to human AD.

## Background

Microglia are the resident macrophages and primary immune effectors in the brain that are central to neuroinflammatory mechanisms of Alzheimer’s disease (AD) pathogenesis [1, 2]. The causal role of microglia in AD is evidenced by data from recent unbiased genome-wide association studies (GWAS) that have identified several single-nucleotide polymorphisms (SNPs) in immune related genes as independent risk factors for late-onset AD [3, 4]. Approximately two thirds of the 29 susceptibility genes are exclusively or most highly expressed in microglia [3, 4]. In human AD and mouse models of AD pathology brains, microglia with activated morphology are frequently found surrounding amyloid-beta (Aβ) plaques [5, 6]. Microglial depletion in mouse models of AD pathology also results in decreased neuropathological features of AD [7, 8]. Overall, these convergent findings implicate microglia in the pathogenesis of AD, but whether microglia are protective [5, 6] or detrimental [1, 2] remains elusive.

Defining the molecular characteristics of microglia in the brain has been primarily undertaken using transcriptomic strategies. Microglial isolation required for these studies can be performed by fluorescence activated cell sorting (FACS), magnetic activated cell sorting (MACS), differential gradient centrifugation, and immunopanning, resulting in collection of intact microglia from fresh mouse and human brain tissues [9–12]. Next generation RNA-sequencing has enabled the quantification of transcripts in low numbers of microglia isolated from brain tissue. Transcriptomic profiles of isolated microglia from animal models have revealed regional heterogeneity of microglia and have shown that progressive neuropathology in AD models results in a shift from homeostatic to transcriptionally distinct disease-related phenotypes [13–18]. Though comprehensive transcriptomic analyses have proven useful in the characterization of microglia, there is still a gap in our understanding of what transcriptional profiles mean biologically and how they relate to protein function. For instance, mRNA and protein expression profiles differ quantitatively, temporally, and spatially [19]. Several studies have shown that transcript-level and protein-level expression data are discordant because of post-transcriptional processes such as mRNA regulation, post-translational protein modifications, and protein recycling and degradation [20, 21]. Protein-mRNA correlation coefficients reach no higher than 0.47 in complex tissues like the brain and this relationship is poorly understood in acutely isolated brain cell types, including microglia [22]. Therefore, global and comprehensive proteomic profiling of isolated microglia can make the most proximate assessment of their biological functions, rather than transcriptional profiling.

Proteomic studies of microglia have primarily used *in-vitro* cultured or neonatal microglia [22–24] and few studies have used CD11b^+^ MACS enrichment to obtain >90% microglia for downstream proteomic analyses. In an effort to elucidate proteomes of adult microglia, we have recently successfully performed quantitative tandem mass tag (TMT) mass spectrometry (MS) of acutely isolated CD11b^+^ MACS-enriched microglia from adult (6-7mo) mice of normal, acute neuroinflammatory (lipopolysaccharide [LPS]-treatment), and chronic neurodegenerative (5xFAD model of AD) states [25]. This deep and comprehensive proteomic study of isolated mouse microglia identified over 4,000 proteins and allowed us to identify unique microglial proteomic changes in mouse models of AD pathology. This study has revealed novel roles for microglial proteins in human neurodegeneration and emphasizes the value of applying state-of-the-art proteomics methods to resolve cell-type specific contributions to disease [25]. Despite microglial enrichment, these data also suggested that MACS still suffers from contamination by proteins from other cell types, most likely due to non-cellular debris that remain in MACS microglial preparations. In support of this limitation, we found that despite >90% microglial cell enrichment by MACS, some of the proteins with highest abundance included oligodendrocytic (Mbp) and astrocytic (Gfap) proteins. Therefore, optimization of microglia purification strategies that are best suited for microglial proteomic studies are warranted prior to comprehensive characterization in disease models as well as in human tissues.

While both MACS-enriched and FACS-based cell purification yield high cell purity, protein-rich non-cellular elements can be definitively excluded by FACS and not by MACS. In this study, we compared the proteomes of MACS-enriched and FACS-isolated CD11b^+^ microglia from adult mouse brain to identify a consensus set of highly-abundant microglial proteins and characterize and contrast the purification efficiencies of each technique. Based on the observed enrichment of microglial proteins and depletion of neuronal, astrocytic, and oligodendroglial proteins, we show that the FACS-isolation approach is the preferred approach for microglial proteomic studies. Using the consensus list of highly-abundant microglial proteins, we validated microglial specific expression of two proteins, namely Msn and Cotl1, in mouse and human brain. We performed *in-vitro* functional studies confirming a role for Msn in Aβ phagocytosis and pro-inflammatory stimulus-induced cytokine release. Importantly, we found Msn as a key member of a glial and inflammatory group of co-expressed proteins that is strongly associated with AD neuropathologies and cognitive dysfunction in human AD cases.

## Methods

### Animals

Mice were housed in the Department of Animal Resources at Emory University under a 12-hour light/12-hour dark cycle with ad libitum access to food and water. All procedures were approved by the Institutional Animal Care and Use Committee of Emory University and were in strict accordance with the National Institute of Health’s “Guide for the Care and Use of Laboratory Animals.”

### Acute isolation of CD11b-positive microglia

The isolation workflow is shown in Figure 1A. Male 4-month-old C57Bl/6J mice (*N* = 10) were anesthetized with isoflurane, followed by exsanguination and cardiac perfusion with 30mL of ice-cold 1× phosphate buffered saline (1×PBS). The brain was immediately dissected and mechanically dissociated over a 40µm cell strainer. Subsequently, the suspension was centrifuged for 5min at 800×*g* at room temperature (RT), supernatant was carefully decanted, and pellet was resuspended in 6mL of 35% stock isotonic Percoll (SIP) solution diluted with 1×HBSS (SIP: 9 parts 100% Percoll and 1 part 10× HBSS). The cell suspension was transferred to a new 15mL conical and 3mL of 70% SIP was slowly underlaid. The established gradient was centrifuged for 25min at 800×*g* with no brake at 15ºC. The top floating myelin layer was aspirated and 3mL from the 35-70% interphase, containing the mononuclear cells (Figure 1A), was collected without disturbing the 70% layer. The mononuclear cells were washed with 6mL of 1×PBS, centrifuged for 5min at 800×*g*, and cell pellet was resuspended in 100µl of 1×PBS.

**Figure 1.**
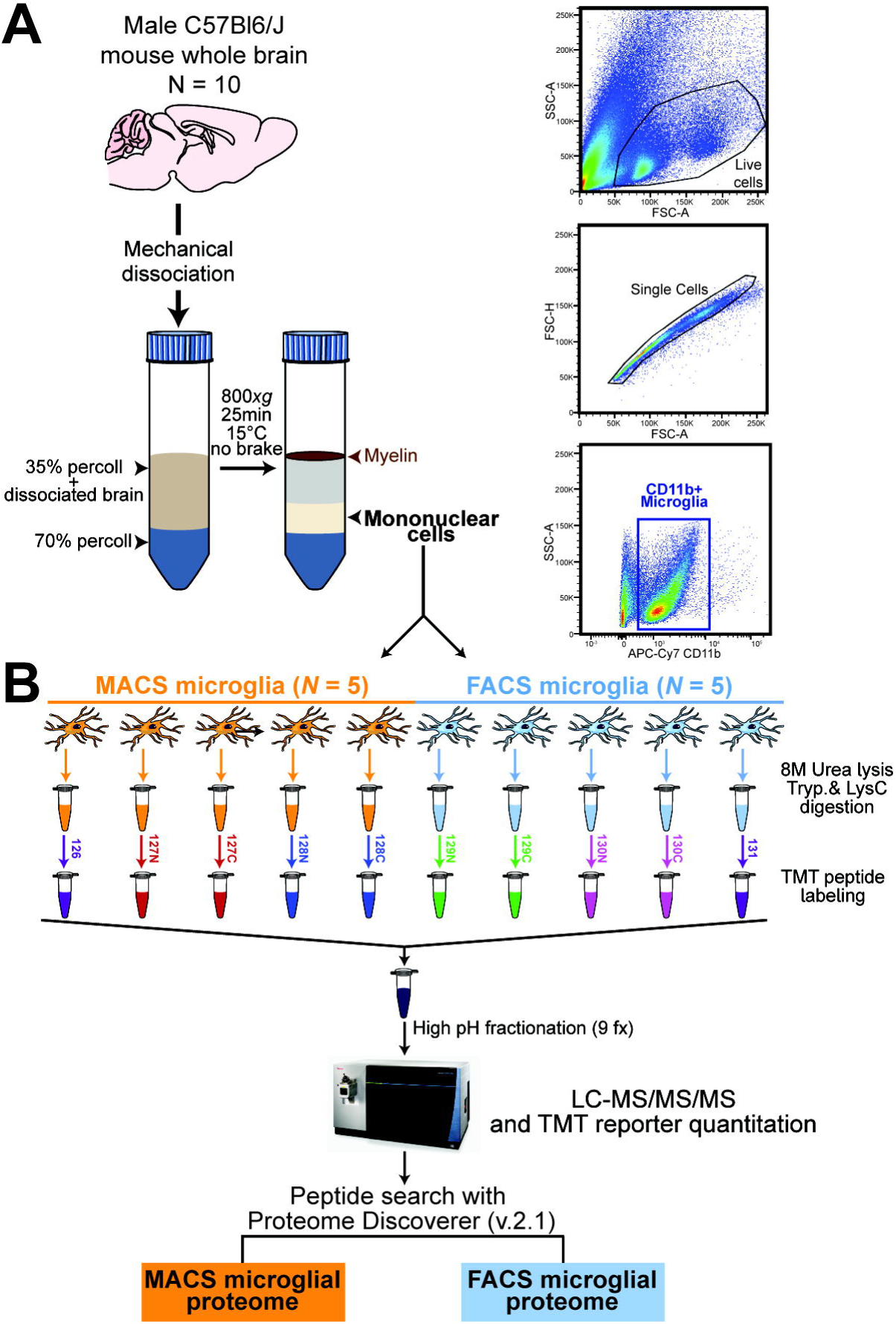
Study design and analytic approach for comprehensive quantitative proteomic analysis of isolated mouse microglia. **A** Workflow summarizing isolation and purification of CD11b^+^ brain immune cells from 4-month-old male C57Bl6/J wild-type mice (*N* = 10). Following mechanical dissociation of fresh, whole mouse brains and percoll density centrifugation, mononuclear cells were enriched for CD11b^+^ microglia cells by magnetic activating cell sorting (MACS, *N* = 5 mice) or isolated via fluorescent activated cell sorting (FACS, *N* = 5 mice). Representative flow cytometry gating strategy and antibody separation using CD11b-APC/Cy7 antibody for isolation of microglia. **B** Proteomic workflow for tandem mass tag (TMT) mass spectrometry (MS) based quantification. All 10 microglia samples were lysed in 8M urea, digested with LysC and Trypsin, and peptides were labeled using one 10-plex TMT kit. A total of 5 individual MACS-enriched microglia samples were dedicated to the first five channels (126, 127N, 127C, 128N, 128C) and 5 individual FACS-isolated microglia samples were dedicated to the last five channels (129N, 129C, 130N, 130C, 131). After labeling, the samples were combined and fractionated by off-line high pH fractionation (*n* = 9 fractions [fx]). Each fraction was analyzed and quantified by synchronous precursor selection (SPS)-MS3 on an Orbitrap Fusion mass spectrometer. Peptide search on raw files were conducted with Proteome Discoverer (v2.1) in order to obtain the MACS microglia proteome and FACS microglia proteome.

Isolated mononuclear cells from five brains were then further purified by CD11b positive selection using MACS columns (*N* = 5 mice, >90% enrichment of CD11b^+^ microglia [25]) (Miltenyi Biotec, Cat. No. 130–093-636) to selectively enrich microglia and brain mononuclear phagocytes (Figure 1B). Cells were pelleted, resuspended in 200µL of lysis buffer (8M urea, 100mM NaHPO_4_, pH 8.5) with HALT protease and phosphatase inhibitor cocktail (ThermoFisher, Cat. No. 78446), and frozen at −80 ºC until protein digestion. Mononuclear cells from the remaining five brains were labeled with APC-Cy7 rat anti-CD11b antibody (BD Pharmingen, Cat. No. 557657) for 30min at RT, washed, and kept on ice until sorting. Compensation experiments were performed using OneComp beads (Thermofisher, Cat. No. 01-1111-42). Live mononuclear cells were gated based on FSC-A/SSC-A, then gated based on FSC-A/FSC-H to identify single cells (Figure 1B), as previously described [26, 27]. The cells were further gated for APC-Cy7 fluorescence, sorted using a BD FACSAria II cell sorter directly into 1.5mL tubes containing 200µL of lysis buffer containing protease and phosphatase inhibitors, and frozen at −80 ºC until protein digestion. Negative controls included unstained mononuclear cells.

In order to allow for a fair comparison between both approaches, we intentionally sorted CD11b^+^ myeloid cells (predominantly microglia) rather than using a CD11b^+^CD45^int^ selection strategy. Since live/dead gating is not feasible for MACS-enrichment, we intentionally did not use live/dead exclusion in the gating strategy during FACS. However, we confirmed in independent experiments that cell viability using our FACS strategy is >95% within CD11b^+^ microglia (**Supplemental Figure 1**). We have also previously shown that CD45^high^ brain-infiltrating macrophages represent <5% of cells within CD11b^+^ brain myeloid cells [25], and therefore, MACS-enriched or FACS-isolated CD11b^+^ brain myeloid cells are referred to as microglia in this manuscript.

### Protein digestion and TMT labeling

Isolated microglia in lysis buffer were sonicated for 3 cycles consisting of 5 seconds of active sonication at 30% amplitude followed by 15 seconds on ice. Protein concentration was determined by bicinchoninic acid (BCA) assay (Pierce, Cat. No. 23225). Protein digestion was performed as previously described [28]. Briefly, 6µg of protein for each sample was reduced with 1mM dithiothreitol (DTT) at room temperature for 30 min and alkylated by 5mM iodoacetamide (IAA) in the dark for 30 min. Samples were then diluted (8-fold) with 50mM triethylammonium bicarbonate (TEAB), digested overnight with Lysyl endopeptidase (Wako, Cat. No. 127-06621) at 1:100 (w/w). The peptide solutions were acidified to a final concentration of 1% formic acid (FA) and 0.1% triflouroacetic acid (TFA), desalted with a C_18_ Sep-Pak column (Waters, Cat. No. WAT054945) and dried down in a vacuum centrifuge (SpeedVac Vacuum Concentrator). TMT labeling of peptides was performed according to manufacturer’s instructions and as previously described [28]. One batch of 10-plex TMT kit (Thermo Fisher, Cat. No. 90110) was used to label all ten samples (Figure 1B). All ten channels were then combined and dried in a vacuum centrifuge.

### High pH reversed-phase peptide fractionation

High pH reversed-phase peptide fractionation kit was used to perform desalting and fractionation as per manufacturer’s protocol (ThermoFisher, Cat. No. 84868). Briefly, the dried sample containing all ten TMT channels was reconstituted with 300µL of 0.1% TFA and added to conditioned reversed-phase fractionation spin columns containing 20mg of resin in a 1:1 water/DMSO slurry. A sequential gradient of increasing acetonitrile (ACN) concentrations in a high-pH elution solution (0.1% triethylamine) was applied to the columns to elute bound peptides into 9 different fractions collected by centrifugation at 3000×*g*. Each fraction was dried in a vacuum centrifuge and stored at 4 degrees until mass spectrometry.

### Mass spectrometry analysis and TMT data acquisition

Assuming equal distribution of peptide concentration across all 9 fractions, 10µL of loading buffer (0.1% TFA) was added to each fraction and 2µL was separated on a 25 cm long by 75 μm internal diameter fused silica column (New Objective, Woburn, MA) packed in-house with 1.9 μm Reprosil-Pur C18-AQ resin. The LC-MS/MS platform consisted of a Dionex RSLCnano UPLC coupled to an Orbitrap Fusion mass spectrometer with a flex nano-electrospray ion source (ThermoFisher). Sample elution was performed over a 120 min gradient with a constant flow rate of 300nl/min. The gradient consisted of multiple steps starting at 3% and going to 7% Buffer B (0.1% formic acid in ACN) over 5 mins, from 7 to 30% Buffer B over 80 mins, from 30% to 60% Buffer B over 5 mins, from 60% to 99% Buffer B over 2 mins, constant at 99% for 8 mins and immediately back to 1% Buffer B for 20 mins. The mass spectrometer was operated in positive ion mode and utilized the synchronous precursor selection (SPS)-MS3 method for reporter ion quantitation as described [28] with a top speed cycle time of 3 seconds. The full scan range was 400–1500 m/z at a nominal resolution of 120,000 at 200 m/z and automatic gain control (AGC) set to 4×10^5^. Tandem MS/MS Collision-induced dissociation (CID) spectra were collected in the ion trap with normalized collision energy set to 35%, max injection time set to 35 ms and AGC set to 1×10^4^. Higher energy collision dissociation (HCD) SPS-MS3 of the top 10 product ions at 65% normalized collision energy (CE) were collected in the orbitrap with a resolution of 60,000, a max injection time of 100 ms and an AGC setting of 5×10^4^.

### Protein identification and quantification

Raw files from Orbitrap Fusion were processed using Proteome Discoverer (v2.1) and MS/MS spectra were searched against UniProt Mouse proteome database (54,489 total sequences) as previously reported [25]. SEQUEST parameters were specified as: trypsin enzyme, two missed cleavages allowed, minimum peptide length of 6, TMT tags on lysine residues and peptide N-termini (+229.162932 Da) and carbamidomethylation of cysteine residues (+57.02146 Da) as fixed modifications, oxidation of methionine residues (+15.99492 Da), and deamidation of asparagine and glutamine (+0.984 Da) as a variable modification, precursor mass tolerance of 20 ppm, and a fragment mass tolerance of 0.6 Da. Peptide spectral match (PSM) error rates were determined using the target-decoy strategy coupled to Percolator [29] modeling of true and false matches. Reporter ions were quantified from MS3 scans using an integration tolerance of 20 ppm with the most confident centroid setting. An MS2 spectral assignment false discovery rate (FDR) of less than 1% was achieved by applying the target-decoy strategy. Following spectral assignment, peptides were assembled into proteins and were further filtered based on the combined probabilities of their constituent peptides to a final FDR of 1%. In cases of redundancy, shared peptides were assigned to the protein sequence with the most matching peptides, thus adhering to principles of parsimony. The search results and TMT quantification as well as raw LC-MS/MS files are included in the ProteomeXchange online repository with identifier PXD015652. We included proteins with TMT abundance values in at least 3 of the 5 replicates per group, as well as proteins present within all 5 replicates in one group and absent in the other group (present only in MACS: total *n* = 4), but also display a high protein FDR confidence (*n* = 2, Gpm6b & Sh3gl2). Additionally, even though TMT labeling limits missing values, normalized abundances of zero were imputed as the lowest TMT abundance value in the dataset.

### Differential expression analysis

Differentially enriched or depleted proteins (unadjusted *p ≤* 0.05) were identified by unpaired t-test comparing the five FACS-isolated microglia samples and the five MACS-enriched microglia samples. Differential expression (FACS/MACS) is presented as volcano plots which display all proteins that either arise from expression of one of the proteins or gene products in the cell type-specific enrichment lists [11, 22]. Significance of differentially expressed protein overlap with cell type-specific enriched protein marker lists was assessed using a one tailed Fisher’s exact test and corrected for multiple comparisons by the FDR (Benjamini-Hochberg) method (**Supplemental Table 1**).

### Gene ontology enrichment analysis

Gene Ontology (GO) functional annotation of differentially expressed proteins was performed using GO-Elite v1.2.5 as previously published [30, 31], with a minimum of five genes per ontology, meeting Fisher exact significance of *p* < 0.05, i.e., a Z-score greater than 1.96. The background gene list for GO-Elite consisted of unique gene symbols for all proteins identified and quantified in our dataset (*n* = 1791). Input lists included proteins that were significantly differentially enriched or depleted (*p* < 0.05 by unpaired t-test, unadjusted) and with a ≥2-fold-change in abundance comparing FACS-isolated with MACS-enriched microglial proteomes (**Supplemental Table 2**).

### Cell-type enrichment analysis

Cell-type enrichment was performed by cross-referencing the number of differentially enriched proteins in our FACS vs. MACS microglial proteomes with cell type-specific gene lists from mass-spectrometry based proteomics [22] (termed “reference cell-type proteome”) and RNA-Seq [11] (termed “reference cell-type transcriptome) of the following isolated mouse brain cell types: microglia, oligodendrocytes, astrocytes, neurons, and endothelial cells. Methods for determining cell-type enrichment of each protein or gene have been previously published and used in prior proteomic analyses [30, 32]. Of note, purified mouse endothelial proteomes have yet to be published.

### Immunofluorescence staining

Brains were isolated from wild-type Cx3cr1^CreER-YFP^ mice (*N* = 4; Jackson Stock No. 021160) and Cx3cr1^CreER-YFP^ mice crossed with 5xFAD mice (*N* = 6) at 6-7 months of age. Cx3cr1^CreER-YFP^ mice constitutively express YFP in microglia [33, 34]. Brains sections from 9-10 month old wild-type (C57Bl6/J, *N* = 3) and 5xFAD (*N* = 4) were used for CD31/Msn and GFAP/Msn co-immunostaining. Briefly, the mice were anesthetized with isoflurane and perfused transcardially with 30mL of 1×PBS. The brains were immediately removed and hemisected along the sagittal midline. The left hemisphere was immersion-fixed in 4% paraformaldehyde for 24 hours, washed 3 times with 1×PBS, and transferred to 30% sucrose for another 24 hours. Subsequently, the brains were cut into 30µm thick sagittal sections using a cryostat. For immunofluorescence staining, 3-4 brain sections from each mouse were thoroughly washed to remove cryopreservative, blocked in 8% normal horse serum diluted in 1× tris buffered saline (TBS) and 0.1% Triton-X for 1 hour, and incubated with primary antibodies diluted in 1×PBS overnight (1:200 goat anti-GFP [Rockland, Cat. No. 600-101-215], 1:100 rabbit anti-Msn [Abcam, Cat. No. ab52490], 1:25 rat anti-CD31 [BD Bioscience, Cat. No. 550274]. 1:500 mouse anti-GFAP [Thermo, Cat. No. 14-9892-82], 1:100 rabbit anti-Cotl1 [Sigma-Aldrich, Cat. No. HPA008918]). For Aβ staining, 5xFAD brain sections were treated with 80% formalin for 3 min, washed with for 10 min per wash, blocked, and incubated overnight with mouse anti-β-amyloid 4G8 clone (1:500, [BioLegend, Cat. No. 800701]). Following washes and incubation in the appropriate fluorophore-conjugated secondary antibody (1:500, anti-goat FITC, anti-rat Alexa 488, anti-mouse FITC, anti-mouse DyLight 405, anti-rabbit Rhodamine-red) for 1 hour, sections were mounted on slides with mounting media containing DAPI (Sigma-Aldrich, Cat. No. F6057) for nuclear staining. Representative images of the same region in the cortex were taken using the Leica SP8 multi-photon confocal microscope and all images processing was performed using Fiji software [35].

### Quantitative analysis of Msn immunofluorescence

Confocal micrographs (40x magnification) from age-matched WT (*N* = 3) and 5xFAD (*N* = 3) brains were used for quantitative analysis of Msn protein expression in microglia. We sampled ramified microglia from WT mice and plaque-associated microglia from 5xFAD mice. Regions of interest containing a single microglial cell were drawn and Msn immunoreactivity per microglial cell was measured by sampling 4-7 microglia per section from each mouse. Background subtraction was performed before analysis. Average Msn immunoreactivity (intensity normalized to area of region of interest) in WT and 5xFAD microglia was statistically compared using unpaired t-test. Image analysis was performed using ImageJ.

### Primary mouse microglia isolation

Primary mouse microglial cultures were established using published isolation and enrichment protocols [36, 37]. Briefly, brains from wild-type C57Bl6/J mice (age P0-3) were removed immediately after euthanization and digested with Trypsin (ThermoFisher, Cat. No. 25300054) for 15 min at 37°C. Subsequently, Trypsin reaction was quenched with 20mL of DMEM (Dulbecco’s Modified Eagle Medium, ThermoFisher, Cat. No. 10569-010) with 10% fetal bovine serum (FBS) and 1% penicillin streptomycin glutamine (ThermoFisher, and Cat. No. 10378016). The floating myelin debris was removed, and the remaining cell suspension was filtered through a 40μm cell strainer and centrifuged at 800×*g* for 5 min. The cell pellet was washed with DMEM/10% FBS followed by CD11b positive selection (Miltenyi Biotec, Cat. No. 130–093-636) using mini-MACS (Miltenyi Biotec, Cat. No. 130-042-201) columns. The CD11b positive microglia were seeded in poly-L-lysine-coated wells and cultured in DMEM at 37°C, 5% CO_2_. The medium was replaced with fresh medium after 24 hours (Figure 4A).

### Small interfering RNA (siRNA) transfection studies

Primary microglia were transfected with 40nM (final concentration) of either Msn siRNA (Santa Cruz Biotechnology, Cat. No. sc-35956) or nonspecific sham siRNA (Santa Cruz Biotechnology, Cat. No. sc-37007) using Lipofectamine™ RNAiMAX (Invitrogen, Cat. No. 13778100) and Opti-MEM (Invitrogen, Cat. No. 31985-062). After 48 hours, the efficiency of siRNA-mediated gene silencing was confirmed by quantitative reverse-transcriptase polymerase chain reaction (qRT-PCR) (Figure 4B).

### Quantitative reverse transcriptase PCR (qRT‐PCR)

Total RNA from microglia was extracted using Trizol (Invitrogen, Cat. No. 15596026) and an RNeasy mini extraction kit (Qiagen, Cat. No. 74104) according to the manufacturer’s instructions. RNA concentration was assessed using Nanodrop and cDNA was synthesized using high-capacity cDNA reverse transcription kit (Applied Biosystems, Cat. No. 4368814). Quantitative real-time PCR was performed on a 7,500 Fast Real-time PCR System (Applied Biosystems) using cDNA, TaqMan PCR Master Mix (Applied Biosystems, Cat. No. 4304437), and gene-specific TaqMan probes (Applied Biosystems) against Moesin (Mm00447889_m1) and Gapdh (Mm99999915_g1). Each primer set was run in triplicate for each RNA sample. Gene expression was normalized to the internal housekeeping gene GAPDH in primary microglial cells, and relative expression was calculated for each gene using the 2ΔΔCT method after normalizing to the control sample (sham siRNA) [38].

### Primary microglia activation by lipopolysaccharide

Microglia were activated with 2ng/mL of lipopolysaccharide (LPS; Sigma Aldrich, Cat. No. L4391, Escherichia coli 0111:B4) after 24 hours of siRNA exposure. Cells were collected after 24 hours of LPS treatment for phagocytosis assays and the supernatants were collected for cytokine assays (Figure 4A).

### Fluorescent fibrillar Aβ42 phagocytosis flow cytometric assay

After *in-vitro* exposure to sham or Msn siRNA and/or LPS stimulus, primary microglia were treated with 2μM (final concentration) of fibrillar fluorescent Aβ42 conjugated to HiLyte Fluor 488 (fAβ42-488, AnaSpec, Cat. No. AS-60479-01) for 1 hour at 37°C. fAβ42-488 was prepared by combining 100μg of peptide with 20μL of 1% ammonium hydroxide (NH_4_OH) and immediately diluted to a final concentration of 100μM with 1×PBS. After incubation at room temperature for 6 days, fAβ42-488 was used for the phagocytosis assay [26, 27]. Subsequent to incubation, cells were treated with trypsin, washed with 1×PBS, and labeled with CD45-PE-Cy7 antibody (BD Biosciences, Cat. No. 552848). Compensation experiments and gating were performed as described above and previously [26, 27]. Single live cells were gated based on CD45-PE-Cy7 fluorescence and fAβ42-488 positivity to determine the phagocytic uptake of Aβ42 within live CD45^+^ microglia (expressed as a proportion of microglia that were positive for Aβ42 fluorescence). Negative controls included microglia incubated with antibodies except for fAβ42-488. We have previously demonstrated loss of fAβ42-488 uptake by microglia after cytochalasin D treatment, confirming that this assay measures actin-dependent phagocytic processes [26, 27].

### Cytokine and chemokine multiplex assay

After *in-vitro* exposure to siRNA and/or LPS stimulus, primary microglia culture supernatants were collected, centrifuged to remove debris, aliquoted, and stored at –80°C. For cytokine analyses, one aliquot of culture supernatant was diluted to 24% in Milliplex Assay Buffer and analyzed using a Milliplex MAP Mouse Cytokine/Chemokine Multiplex Assay (Millipore Sigma, Cat No. MCYTOMAG-70K) per manufacturer’s instructions for the following custom collection of analytes: GM-CSF, IFN-γ, IL-10, IL-1α, IL-2, IL-4, IL-6, IP-10/CXCL10, MCP-1/CCL2, MIP-1α/CCL3, TNF-α, M-CSF, VEGF-A, G-CSF, RANTES [39]. Fluorescent intensity readings for each cytokine were compared across experimental groups and performed in biological triplicates (Figure 4A).

### Existing transcriptomic and proteomic datasets used for comparative analyses

#### Human brain proteomic data

##### Dataset 1

Msn protein abundance, neuropathological, and cognitive data were obtained from a recently published study [40] (Synapse Web Portal 10.7303/syn20933797) and are shown in Figure 5B and 5D. In this study, dorsolateral prefrontal cortex (DLPFC) tissues from non-disease control (*N* = 91), asymptomatic AD (AsymAD, *N* = 98), and AD (*N* = 230) cases were analyzed by mass spectrometry based proteomics using label-free quantitation (LFQ). A total of 3,334 proteins were quantified with fewer than 50% missing values across all cases and were used to make a protein co-expression network using the weighted correlation network analysis (WGCNA) algorithm. This dataset was selected because it is currently the largest quantitative mass spectrometry-based proteomics analysis that generated a consensus AD brain protein co-expression network across multiple study centers.

##### Dataset 2

Msn protein abundance values were obtained from a recently published study [40] (Synapse Web Portal 10.7303/syn20933797) and are shown in Supplemental Figure 6A. In this study, control (*N* = 43), AD (*N* = 47), f rontotemporal dementia with TDP-43 pathology (FTLD-TDP, *N* = 29), amyotrophic lateral sclerosis (ALS, *N* = 54), and Parkinson’s disease and Parkinson’s disease dementia (PD/PDD, *N* = 76) DLPFC brain tissues were analyzed by LFQ-MS and 2,919 proteins were identified. This dataset was selected to determine whether increased Msn levels in AD brain replicated in a different cohort and to see if this increase is selective to AD.

##### Dataset 3

Msn protein abundance values were obtained from a recently published study (Synapse Web Portal 10.7303/syn20933797) [40] and are shown in Supplemental Figure 6B. Control (*N* = 13), AsymAD (*N* = 13), and AD (*N* = 22) brain tissue from precuneus were analyzed using LFQ-MS and 3,348 proteins were identified. This dataset was selected to assess whether Msn levels increase in another brain region in addition to the DLPFC from Dataset 1 and because the precuneus is a region affected early in AD pathogenesis.

##### Dataset 4

Msn protein abundance data were obtained from a recently published study by Drummond *et al.* [41] and are shown in Supplemental Figure 6C. In this study, LFQ-MS was performed on Aβ plaques microdissected from the hippocampus and adjacent entorhinal cortex of rapidly progressive AD (rpAD, *N* = 22) and sporadic AD (spAD, *N* = 22) cases. Approximately 900 proteins were quantified from every case. This is the first large-scale proteomic study that analyzed the Aβ plaque proteome and thus, was used to validate our observation of Msn positive microglia surrounding Aβ plaques in the 5xFAD mouse brain.

#### Human brain transcriptomic data

Msn gene expression in pre-frontal cortex from control (*N* = 157) and AD (*N* = 308) brains was obtained from Zhang *et al.* (GEO: GSE44772) [42]. For each sample, 39,579 transcripts were profiled which represent 25,242 known and 14,337 predicted gene-expression traits. The gene-expression was adjusted for age and sex, postmortem interval (PMI) in hours, and sample pH and RNA integrity number (RIN).

#### Mouse brain proteomic data

We obtained Msn abundance from an existing proteomic dataset [43] (PXD006214) of WT, 5xFAD (Aβ pathology), JNPL3 (tau pathology), and 5xFAD crossed with JNPL3 (Aβ and tau pathologies) mouse lines. In this study, quantitative mass spectrometry using TMT tags was performed on hippocampi from mice of all four lines at 4 months, 7 months, and 10 months of age with three biological replicates for each line and age. The samples were randomized into four 10-plex TMT batches and a total of 6,964 proteins were identified after statistical processing. This dataset was selected because it provides a deep proteome of two mouse models of AD pathology, Aβ and tau, at various ages.

#### Mouse brain transcriptomic data

Msn gene expression in cortical tissue from WT and 5xFAD mice at 3 months, 6 months, and 12 months of age was obtained from Bai *et al.* [44] (GEO: GSE140286). There were 2-3 animals per genotype and age. The gene-level fragments per kilobase of transcript per million (FPKM) values were computed from the read counts for each gene.

### Immunostaining of human brain tissues

Cryopreserved, pathology-confirmed AD (*N* = 10, 5 males, 5 females, average age at death = 72.7) and non-disease control (*N* = 10, 5 males, 5 females, average age at death = 61.4) frontal cortex tissue sections (50μm thick) were obtained from the Emory Neuropathology/Histochemistry Core. Sections were immunostained for Msn (1:100, Abcam, Cat. No. ab52490) using DAB as the chromophore. Adjacent sections were processed for double fluorescent immunolabeling for Msn and Aβ beta protein (4G8, 1:1000, Signet, Cat. No. 9220-02). Sections were incubated in primary antibody for 72 hours followed by a 2 hour incubation in secondary antibodies conjugated to appropriate fluorophores. Secondary antibodies used were Alexa 488 goat anti-mouse/rabbit (1:200, Molecular Probes, Eugene, OR) and Cy3 or biotinylated goat anti-mouse/rabbit (1:200, Jackson Immunoresearch Labs). Subsequently, IF sections were incubated in diamidino-2-phenylindole (DAPI) for 10min (1:5000, Molecular Inc Probes, Eugene, OR). For controls primary antibodies were (a) either pre-absorbed with specific peptide sequence or (b) were omitted. Autofluorescence was eliminated (Chemicon, Cat. No. 2160), sections were mounted, and coverslipped. Images were captured using an Olympus bright-field and fluorescence microscope and camera (OlympusBX51). For final output, images were processed using Adobe Photoshop software.

### Statistical considerations

Our statistical approach for MS studies is described above. One-way analysis of variance (ANOVA) was used to identify group-wise differences and post-hoc Tukey’s test was used to identify pairwise differences (*p* < 0.05 considered significant). All analyses were performed using Graphpad Prism (v7).

## Results

### Comparative proteomic analysis of MACS and FACS-based strategies reveals highly-abundant core microglial proteins and highlights differences between both purification strategies

We applied TMT-MS to obtain the proteomes of CD11b^+^ microglia from 4-month-old male C57Bl6/J wild-type mice (*N* = 10) isolated using two different strategies: magnetic-activated cell sorting (MACS *N* = 5, >90% enrichment [25]) and fluorescence activated cell sorting (FACS, *N* = 5, average 30,000 total cells per sample). MACS-enriched and FACS-isolated microglia cell lysates in 8M urea were digested with LysC and Trypsin, labeled with isobaric multiplex TMTs, and analyzed by synchronous precursor selection (SPS)-based MS3 (SPS-MS3) (Figure 1B). Due to the isobaric nature of the tags, all shared peptides from the 10 samples exhibit the same biochemical properties (i.e., exact mass, ionization efficiency, and retention time) in the MS1 or precursor scan. Only following MS/MS (or MS3-SPS) does each tag fragment and release unique reporter ions, which are then used for peptide quantitation. Thus, one major advantage of multiplex TMT based quantification is that the peptides are pooled across all samples, which increases the precursor peptide signal in the mass spectrometer for proteins common to all samples by up to 10-fold (e.g. TMT 10-plex) when compared to running each sample individually by LFQ. This is especially important when isolating small numbers of cells (~30,000) as in this study. In total, we identified 6,404 peptides mapping to 1,876 protein groups. Of these, 1,791 proteins were quantified in at least 3 of the 5 replicates in each group, and 4 demonstrated expression patterns unique to only the MACS-enriched group, of which we excluded 2 due to low protein FDR. Despite the potential relative advantages or disadvantages of FACS and MACS-based microglial purification strategies, we hypothesized that core microglial proteins must be measured with high abundance by both approaches. We first focused on consensus between both the proteomes and identified 10 proteins (Hmga1, Msn, Pfn1, Myh9, Coro1a, S100a9, Cap1, Cotl1, Ctsd) that satisfied the following criteria: (1) highly abundant (>90^th^ percentile) in MACS and FACS microglial proteomes in our data, and (2) most highly expressed at the protein level in microglia as compared to neurons and other glial cells based on an existing cell type-specific proteomic reference database [22].

Although FACS and MACS based purification of microglia yield high microglial purity (>90% for both), significant differences in the types of proteins identified may exist. Comparing FACS and MACS microglial proteomes, a total of 953 proteins were differentially enriched (*p* < 0.05, unpaired t-test), and of these, 815 met significance thresholds at the FDR <5% level. Of the 953 differentially enriched proteins, 685 were increased and 268 proteins were decreased in abundance in the FACS microglia proteome. When we compared the relative abundance of the 685 significantly increased proteins, 36 proteins had a 2-fold or greater increase in abundance (Figure 2A, solid orange dots delineated by dotted line) in the FACS proteome than in the MACS proteome (top 5: Apex1, Fam3c, Lcn2, Lbr, S100a9). Of the 268 significantly decreased proteins, 65 proteins had a 2-fold or greater decrease in abundance in the FACS proteome (Figure 2A, solid blue dots delineated by dotted line), i.e., an increased abundance in MACS proteome (top 5: Gap43, Gpm6b, Sh3gl2, Psip1, Hmgn3). Gene ontology (GO) analysis of proteins significantly decreased (*n* = 268) in the FACS microglia proteome indicated an over-representation of “mitochondrial” and “synaptic” proteins, and proteins involved in “electron transport chain”, “neurotransmitter transport”, and “synaptic transmission” processes (Figure 2B). Conversely, “cytosolic”, “endoplasmic reticulum (ER)”, and “ribosome” cellular components, and proteins involved in processes such as “regulation of protein metabolism” and “immune system” were overrepresented by the proteins significantly increased (*n* = 685) in the FACS microglia proteome (Figure 2B). The cytosolic and ER bias of the FACS proteome, argues against a nuclear proteomic bias that could be expected using the FACS approach.

**Figure 2.**
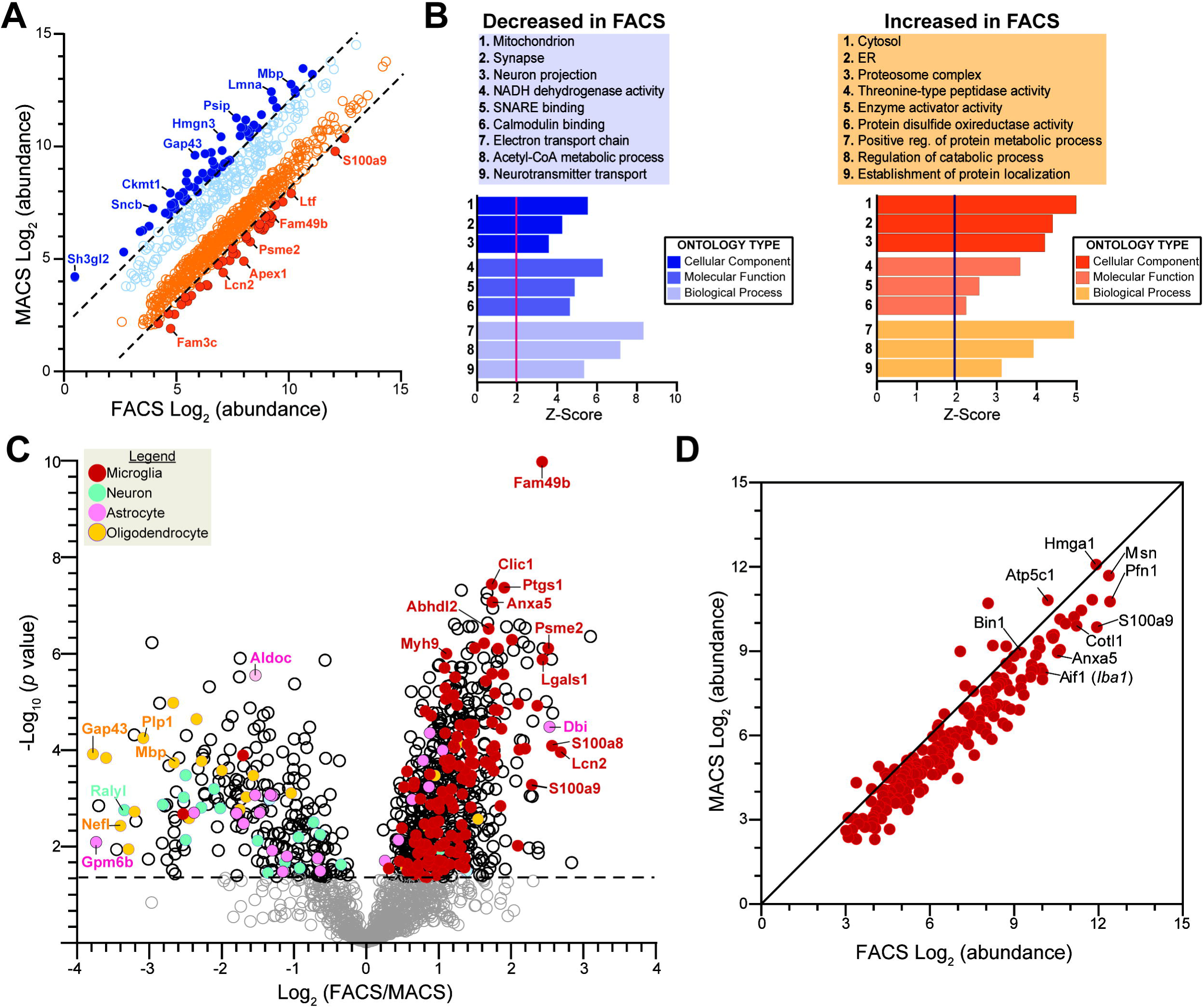
Enrichment of microglia-specific proteins and depletion of non-microglial proteins by FACS. **A** Scatter plot displaying relative abundance of all differentially expressed proteins (*n* = 953) between FACS-isolated microglia proteome and MACS-enriched microglia proteome (all dots represent *p* < 0.05). Dotted lines differentiate proteins with a Log_2_ fold-change of at least 2. Solid orange dots indicate proteins significantly higher in the FACS proteome or lower in MACS proteome. Solid blue dots indicate proteins significantly higher in MACS proteome or lower in FACS proteome. **B** Results from Gene Ontology (GO) enrichment analysis of 953 differentially expressed proteins against a background list of total proteins identified in current dataset. Representative GO terms (top 3) from each of the three GO groups. Degree of enrichment of each GO term is indicated by the Z-score (X-axis). Vertical red and black lines indicate a Z-score of 1.96. **C** Volcano plot displaying the distribution of differentially expressed proteins between FACS-isolated and MACS-enriched microglia proteomes. Cell type-specific markers defined by a reference proteome, Sharma et al. [22], shows significant enrichment of microglial specific proteins in the FACS proteome (*p* < 0.05, Unpaired t-test). Red dots = microglia, turquoise dots = neuron, pink dots = astrocyte, yellow dots = oligodendrocyte. Grey dots represent differentially expressed proteins with a *p* > 0.05. Log_2_ fold-change is shown on the X-axis, -Log_10_(*p*-value) is shown on the Y-axis, and horizontal dotted line indicates *p* value of 0.05. **D** Consensus microglial proteins between MACS and FACS microglial proteomes (*n* = 203). From this list, Moesin (Msn) was chosen for further validation.

These results show that despite using identical mass spectrometry approaches and high cellular purity obtained by MACS and FACS-based microglial enrichment approaches, the proteomes from each strategy are indeed very different. Namely, the proteome of MACS-enriched microglia appears to be heavily biased towards synaptic proteins, suggesting a significant neuronal component in these samples, while the FACS-isolated proteome is enriched for immune function, indicating higher enrichment of microglial proteins. Despite these methodological differences, we identify a core set of highly-abundant microglial proteins (Figure 2D, n = 203).

### FACS-isolated microglial proteomes result in higher enrichment of microglial proteins and depletion of non-microglial protein contaminants

As compared to MACS, FACS has the added advantage of confidently gating out non-cellular but protein-rich elements present in cell preparations from the brain. Based on this and our observed proteomic differences between FACS and MACS-based approaches, we hypothesized that the FACS-based strategy results in greater enrichment of microglial proteins and depletion of non-microglial proteins that may contaminate mononuclear cell preparations from the brain. Therefore, we performed cell-type enrichment analysis of differentially expressed proteins using protein marker lists derived from existing reference proteomes of four purified mouse brain cell-types — microglia, neurons, astrocytes, and oligodendrocytes [22] (Figure 2C). Although endothelial cells represent a key brain cell population, at the time of this work, there was no existing reference for purified endothelial cell proteome. Among the 953 proteins differentially expressed comparing FACS vs MACS proteomes, we observed a marked enrichment of proteins highly expressed by microglia in the FACS proteome. The microglial proteins with high FACS-proteome abundance included Fam49b, Ptgs1, Clic1, Anxa5, and Psme2 (Figure 2C, red solid dots). In general, microglia-specific proteins were quantified at higher abundances in FACS compared to MACS proteomes (Figure 2D, **Supplemental Figure 2A**). Conversely, neuronal, astrocytic and oligodendroglial proteins were relatively highly abundant in the MACS proteome (**Supplemental Figure 2B-D**).

In addition to utilizing a reference brain cell type-specific proteome [22], we further analyzed our data for cell-type enrichment using a reference transcriptomic (RNAseq) dataset of five purified mouse brain cell types— microglia, neurons, astrocytes, oligodendrocytes, and endothelial cells [11] (**Supplemental Figure 2E**). Similar to our observations using reference proteomic data, we observed higher representation of microglia-specific genes among proteins enriched in the FACS proteome while neuronal, astrocytic and oligodendroglial genes were more highly represented in the MACS proteome (**Supplemental Figure 2E**). Interestingly, genes highly abundant in endothelial cells were also more highly abundant in the FACS proteome (**Supplemental Figure 2E**, green dots). This intriguing enrichment of endothelial markers in the FACS-based microglial proteome could indicate molecular overlap between microglia and endothelial cells or could highlight limitations of prior endothelial transcriptomes. However, this can likely be further resolved by future purified endothelial proteomic studies.

Since both reference proteomic and transcriptomic cell type-specific datasets used for our analyses were independent, we assessed the degree of overlap between cell specific markers identified by these references (**Supplemental Figure 2F**). Interestingly, the two reference cell-type marker lists we used identified minimally overlapping and varying numbers of proteins for each cell-type. For example, of the 140 microglia-specific proteins identified at the proteomic level, only 87 were also identified as enriched in microglia at the transcriptomic level (**Supplemental Figure 2F**). Despite these differences in reference datasets, the consensus between both our cell enrichment analyses highlights the validity of our findings regarding the advantages of the FACS approach over MACS enrichment for microglial proteomics. A detailed summary of differential representation of brain cell type-specific genes and proteins is provided in Table 1.

**Table 1.** Summary of cell-type enrichment in differentially expressed proteins between MACS and FACS isolated microglia proteomes.

| Reference | Cell type | Increased in FACS<br>n (%) | Decreased in FACS<br>n (%) | Total<br>n |
| --- | --- | --- | --- | --- |
| **Reference**<br>**cell-type**<br>**proteome**<br>*Sharma et al. 2015* | **No cell-type** | 520 (70.6) | 217 (29.4) | 737 |
|  | **Microglia** | 139 (99.3) | 1 (0.7) | 140 |
|  | **Neuron** | 2 (10) | 18 (90) | 20 |
|  | **Astrocyte** | 19 (57.6) | 14 (42.4) | 33 |
|  | **Oligodendrocyte** | 5 (21.7) | 18 (78.3) | 23 |
| **Reference**<br>**cell-type**<br>**transcriptome**<br>*Zhang et al. 2014* | **No cell-type** | 515 (71.3) | 207 (28.7) | 722 |
|  | **Microglia** | 119 (100) | 0 | 119 |
|  | **Neuron** | 4 (8) | 46 (92.0) | 50 |
|  | **Astrocyte** | 7 (43.8) | 9 (56.2) | 16 |
|  | **Oligodendrocyte** | 11 (73.3) | 4 (26.7) | 15 |
|  | **Endothelial cell** | 29 (93.5) | 2 (6.5) | 31 |

To further confirm the validity of the apparent cellular biases of MACS vs. FACS proteomes in our dataset, we compared our results to our recently published MACS-based mouse microglial proteome from wild-type mice and 5xFAD mouse models in which 4,133 proteins were identified by TMT-MS in CD11b^+^ MACS-enriched microglia [25]. In this secondary analysis of an independent MACS-enriched microglial proteome, we cross-referenced the data with the reference cell-type proteomes [22] and found that 78.5% of all proteins did not map to a given cell-type, while only 4.5% were microglial and 17% of the proteins were enriched in other cell-types (**Supplemental Figure 3A**). While microglia-specific proteins such as Msn and Cotl1 were highly abundant in the MACS-enriched microglial proteome, neuronal proteins (Camk2a, Gap43), astrocyte proteins (Gfap, Aldoa) and oligodendrocyte proteins (Mbp) were also identified as equally highly abundant proteins (**Supplemental Figure 3B**, >90^th^ percentile of relative abundance). Non-microglial proteins were also found in the FACS-isolated microglial proteome at relatively high abundance (**Supplemental Table 1**). For example, Mbp (90^th^ percentile), Gfap (60^th^ percentile), Camk2a (>75^th^ percentile) were still relatively abundant in the FACS microglial proteome, although many fold lower compared to the MACS proteome where these three proteins were found at >90^th^ percentile of abundance.

Collectively, these cell-type enrichment analyses of our proteomic data show that MACS-based proteomes are still contaminated by non-microglial proteins despite high microglial cellular purity likely due to remnant acellular debris; while FACS-based microglial isolation results in a purer microglial proteome that is less contaminated by neuronal, astrocytic and oligodendroglial proteins. The presence of non-microglial proteins at relatively high abundance in the FACS-isolated microglial proteome may be more indicative of phagocytic uptake on non-microglial cellular elements by microglia [45]. Another interesting finding is that a majority of proteins expressed in microglia are not specific to microglia but are shared across various cell types.

### Cotl1 and Msn are highly expressed by microglia in adult mouse brain

Based on our consensus findings between different microglia enrichment strategies, we have defined a core set of highly-abundant microglial proteins (Figure 2D, *n* = 203) in adult mice which can be used as indicators of microglial abundance in future proteomic studies. Among proteins with high concordance in microglia, several proteins including Msn, Cotl1 and Bin1 are identified to be highly abundant in both FACS and MACS proteomes (Figure 2D). We chose Cotl1 and Msn for additional neuropathological studies to characterize expression in the mouse brain. Coactosin like F-actin binding protein 1 (Cotl1) is an actin-binding protein expressed by immune cells including macrophages and has been recently reported to regulate actin dynamics at the immune synapse [46, 47]. Cotl1 along with Msn also bind polyunsaturated fatty acid chemokines [48].. Furthermore, Cotl1 is highly and specifically expressed by microglia at the transcript and protein levels [11, 22]. we have previously identified as a novel microglia-specific marker from a quantitative TMT-MS proteomic analysis of MACS-enriched CD11b^+^ microglia of adult mice in normal, acute neuroinflammatory, and chronic neurodegenerative states [25]. In the current study, we were able to replicate our previous finding of Cotl1 as a microglial marker with a significantly higher abundance (1.4-fold higher, *p* = 0.001) in the FACS proteome than MACS proteome. We immunostained brains from 6-7 month old Cx3cr1^CreER-YFP^ mice on wild-type (WT) or 5xFAD backgrounds with antibodies against YFP/GFP, to detect microglia, and Cotl1. Cx3cr1^CreER-YFP^ mice express YFP/GFP immunofluorescence in Cx3cr1^+^ microglia in the brain [33, 34]. In the Cx3cr1^CreER-YFP^-WT and Cx3cr1^CreER-YFP^-5xFAD cortices, Cotl1 immunofluorescence was specifically detected in GFP immunofluorescent microglia (**Supplemental Figure 4**, arrow). The GFP^+^ microglia in the Cx3cr1^CreER-YFP^-WT cortex also appeared morphological ramified, but those in Cx3cr1^CreER-YFP^-5xFAD cortex adopted an activated morphology with shorter swollen processes and larger cell bodies (**Supplemental Figure 4**). We have previously described a similar morphological phenotype in a WT and 5xFAD mouse brains as well as demonstrated a quantitative increase of Cotl1 expression in 5xFAD brain compared to WT brain [25].

Moesin (Msn) is a member of the ezrin-radixin-moesin (ERM) family of proteins that link the C-terminal domain of cortical actin filaments to the plasma membrane [49]. Outside the brain, Msn in found in other tissues and cell types including macrophages, lymphocytes, epithelial cells, and endothelial cells [50–52]. In the brain, Msn is most abundantly expressed by microglia compared to neurons, oligodendrocytes or astrocytes although a brain endothelial cell proteome is currently lacking [22]. At the transcript level, Msn is highly expressed by endothelial cells and microglia [10, 11, 22]. We immunostained the brains from Cx3cr1^CreER-YFP^ mice on WT or 5xFAD backgrounds for YFP/GFP (to detect microglia) and Msn. In Cx3cr1^CreER-YFP^-WT mouse cortex, we observed GFP immunofluorescence within the cell body and processes of ramified microglia (Figure 3A, arrow) and detected diffuse Msn immunofluorescence in the same ramified microglia (Figure 3A, arrow). We also observed Msn immunofluorescence in non-microglial cells which resembled endothelial cells (Figure 3A, asterisk), consistent with previously reported Msn expression in endothelial cells [11]. In Cx3cr1^CreER-YFP^-5xFAD mouse cortex, we observed GFP immunofluorescent microglia with an activated morphology and found strong Msn expression in the same microglia (Figure 3A, arrow). We also observed GFP and Msn immunofluorescent microglia surrounding dense Aβ plaques (Figure 3B, arrow) in Cx3cr1^CreER-YFP^-5xFAD mouse cortex. Importantly, 5xFAD mice displayed a significant increase in Msn immunofluorescence in the cortex compared to control mice (**Supplemental Figure 4B**, *p* < 0.0001). Given that Msn is abundant in endothelial cells at the transcript level and the presence of GFP negative, Msn immunofluorescent non-microglial cells (Figure 3A, asterisk), we stained brain tissue from 9-10 month old WT and 5xFAD mouse brains to detect CD31 (an endothelial marker) and Msn. In both WT and 5xFAD cortices, we observed CD31 and Msn immunofluorescence in endothelial cells (Figure 3C, arrows) as well as cells immunofluorescent for Msn only (Figure 3C, asterisk). Additionally, we stained the same brain sections with GFAP, to detect astrocytes, and Msn. In WT and 5xFAD cortices, GFAP immunofluorescent astrocytes (Figure 3D, asterisk) did not display Msn immunofluorescence (Figure 3D, arrow). Overall, while Msn is found in both ramified microglia and endothelial cells in normal brain and in regions distant from Aβ plaques, Msn expression is nearly exclusively found in microglia that surround Aβ plaques in 5xFAD mice.

**Figure 3.**
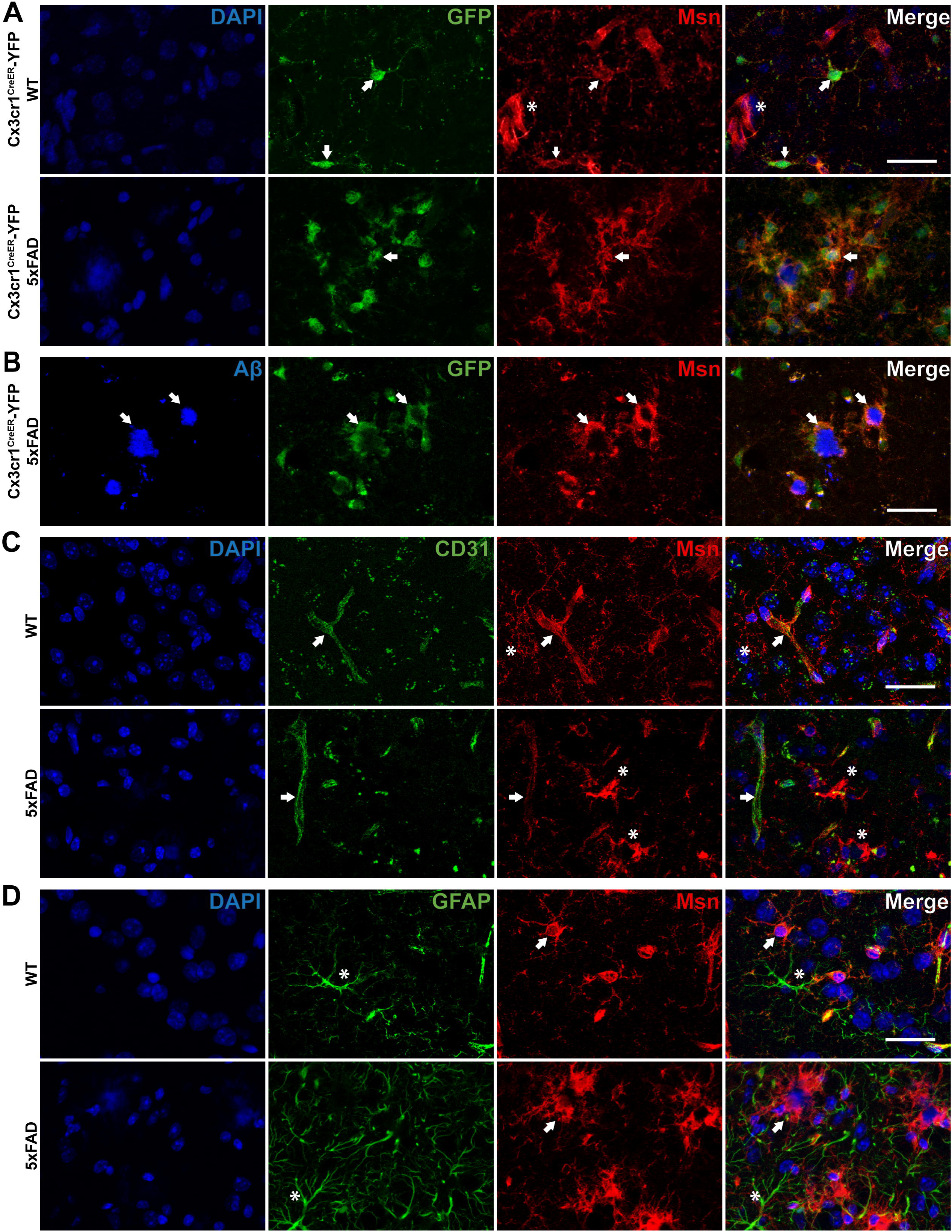
Moesin is expressed by microglia and endothelial cells in mouse brain. **A** Representative immunofluorescence images of Cx3cr1^CreER-YFP^-WT (*N* = 4) and Cx3cr1^CreER-YFP^-5xFAD (*N* = 6) mouse cortex stained for microglia (GFP) and Msn. Arrow indicates microglia immunopositive for GFP and Msn. Asterisk indicates cells immunopositive for Msn only. **B** Representative immunofluorescence images of Cx3cr1^CreER-YFP^-5xFAD (*N* = 6) mouse cortex stained for amyloid-beta (Aβ), microglia (GFP), and Msn. Arrow indicates Aβ plaque as well as microglia immunopositive for GFP and Msn. **C** Representative immunofluorescence images of WT (*N* = 3) and 5xFAD (*N* = 4) mouse cortex stained for endothelial cells (CD31) and Msn. Arrow indicates endothelial cells immunopositive for CD31 and Msn. Asterisk indicates cells immunopostive for Msn only. **D** Representative immunofluorescence images of WT (*N* = 3) and 5xFAD (*N* = 4) mouse cortex stained for astrocytes (GFAP) and Msn. Arrow indicates cells immunopositive for Msn only. Asterisk indicates astroctyes immunopostive for GFAP only. Scale bar = 30µm.

These results validate two highly-abundant microglial specific proteins, namely Cotl1 and Msn, which can serve as markers of microglia in the mouse brain at normal and disease states and indicate possible roles for these proteins in microglial function under homeostatic and disease conditions.

### Msn regulates Aβ phagocytosis and pro-inflammatory cytokine production by microglia

Based on our findings of Msn expression in mouse microglia distant and proximate to Aβ plaques in the 5xFAD mouse brain, we performed *in-vitro* functional studies using primary mouse microglia to determine the effect of Msn knockdown (KD) on microglial phagocytic properties (Figure 4A). Msn siRNA resulted in >90% reduction in Msn mRNA expression compared to control siRNA (Figure 4B, *p* < 0.001). Primary neonatal mouse microglia were incubated with fluorescent-conjugated fAβ42 for 60 min and phagocytic uptake by CD45^+^ live microglia was assessed by flow cytometry [26]. Msn siRNA pre-treatment reduced Aβ phagocytic uptake by ~25% (*p* = 0.0274) without affecting viability as compared to siRNA control under resting conditions (Figure 4B). In the presence of LPS stimulation, we observed minimal effect of Msn siRNA (Figure 4B), suggesting context-dependent functional roles for Msn in microglia. In this assay, we have previously shown that inhibition of actin-dependent processes by cytochalasin D completely inhibits Aβ uptake, confirming actin-dependent phagocytic uptake of Aβ rather than passive binding to the cell surface [26]. Next, we measured levels of cytokines and chemokines by Luminex assays in the culture supernatants derived from primary microglia that were treated with sham siRNA or siMsn for 24h and then treated with LPS (or sham) for another 24h as a strong pro-inflammatory challenge (Figure 4A). As expected, LPS increased levels of TNF, IL-10, and other cytokines (Figure 4D-I, IL6, G-GSF, IP-10/CXCL10, and MIP-1α/CCL3). Specifically, Msn siRNA enhanced LPS-induced increase in IL-10 (Figure 4D) and TNF (Figure 4F, *p* < 0.05) levels, although the former was not statistically significant. In sum, our *in-vitro* studies indicate that deletion of Msn from microglia impairs their ability to phagocytose Aβ and increases IL-10 and TNF-α release after an inflammatory stimulus.

**Figure 4.**
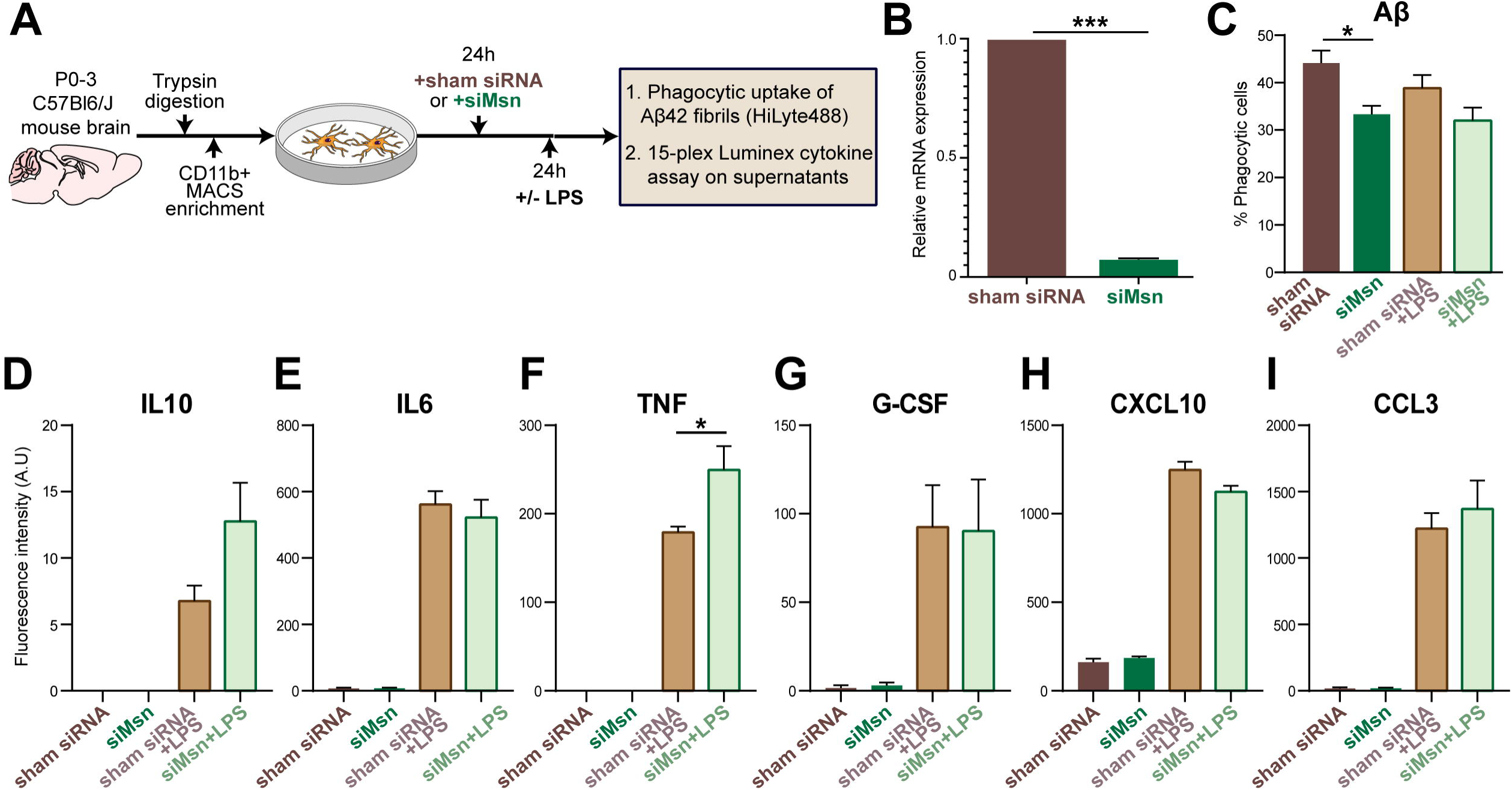
Moesin knockdown impacts Aβ phagocytosis and pro-inflammatory cytokine production by microglia. **A** Overview of *in-vitro* knockdown studies in primary mouse microglia treated with Msn siRNA or sham siRNA under resting and LPS-stimulated conditions. **B** Results from qRT‐PCR experiments demonstrating efficiency of Msn knockdown by siRNA as compared to sham siRNA. Y‐axis represents relative mRNA expression (2‐ΔΔCt method) normalized to Gapdh as housekeeping gene (three independent biological replicate experiments were performed per condition). **C** *In‐vitro* phagocytic capacity of primary microglia for fAβ42 HiLyte488 following exposure to sham siRNA or Msn siRNA under resting and LPS-stimulated conditions. Phagocytic uptake of fAβ42 was measured as proportion of CD45^+^ microglia using untreated microglia as negative controls. For each sample, at least 2,000 live CD45^+^ microglial events were captured. *N* =□3 independently performed biological replicate experiments. **D-I** Bar graphs displaying cytokine/chemokine data, shown as fluorescence intensity in arbitrary units (A.U.), obtained from microglial culture supernatants after siRNA sham or siRNA Msn treatment under resting and LPS-stimulated conditions: **D** IL10, **E** IL6, **F** TNF, **G** G-CSF, **H** CXCL10, **I** CCL3. Error bars represent ± SEM. Unpaired t-test: \**p* <□0.05, \*\*\**p* <□0.0001.

### Msn is a hub protein in a disease-associated proteomic co-expression module in human AD brain

In a recent large proteomic analysis of over 400 human post-mortem brains, we identified a consensus network of protein co-expression modules with robust associations to AD pathologies as well as cognitive traits [40]. In this network analysis, we identified a protein module (M4) that was highly correlated with higher neuropathological burden and worse cognitive outcomes; this module was highly enriched in microglial and astrocytic proteins [40]. Msn is a hub protein, defined as a protein most central to module function with a high module membership value (kME), of this module (Figure 5A, kMe = 0.807) as well as several other microglial proteins identified in the current mouse microglial proteomic analysis (Figure 5A, red circles: ANXA5, CRYL1, TPP1, COTL1, CTSB, and CAP1). Msn protein abundance was significantly elevated in AD DLPFC compared to control or AsymAD DLPFC (Figure 5B, *p* < 0.001). Furthermore, increased Msn levels were also present in AD brains even at the transcript level (Figure 5C, *p* < 0.001) [42] We observed that Msn protein abundance displayed a strong negative correlation with cognitive function (Figure 5D, MMSE, *r* = −0.51, *p* = 1.9e-12). Cognitive function was determined by the mini-mental status examination (MMSE) score where higher scores (>24) represent better cognitive function and lower scores (<24) represent cognitive dysfunction (or dementia) [53]. Furthermore, Msn protein abundance demonstrated a strong positive correlation with Aβ plaque burden (Figure 5D, CERAD, *r* = 0.32, *p* = 2e-11) and neurofibrillary tau tangle pathology (Figure 5D, BRAAK, *r* = 0.33, *p* = 4.2e-12) [54, 55]. We further characterized Msn expression via immunohistochemistry and observed a morphological difference between control and AD brain sections (Figure 5E). Msn immunoreactive cells in the control brain showed a typical, ramified microglial morphology (Figure 5E, Control inset), while Msn immunoreactive cells resembled activated microglia in the AD brain (Figure 5E, AD inset). Lastly, we observed Msn immunofluorescent cells to be surrounding Aβ plaques in AD brain (Figure 5F), similar to what we observed in the 5xFAD mouse brain (Figure 3B).

Using independent proteomic datasets, we examined Msn protein abundance in published human post-mortem brain cohorts which included a variety of neurodegenerative disease cases [40, 41]. Similar to our primary findings (Figure 5B), Msn abundance was significantly increased in AD DLPFC compared to controls in an independent cohort (**Supplemental Figure 5A**, *p* < 0.001). Interestingly, we observed a significant elevation in Msn abundance in FTLD-TDP DLPFC compared to controls (**Supplemental Figure 5A**, *p* < 0.001); however, this elevation was not seen in ALS and PD/PDD brains (**Supplemental Figure 5A**). Next, we wanted to determine whether Msn levels were increased in a different brain region other than DLPFC; thus we analyzed a proteomic analysis of the precuneus in control, AsymAD, and AD cases [40]. We chose this region because the precuneus is affected early in the course of AD, as shown by clinical, imaging, and pathological studies[56]. We found Msn to be significantly elevated in AD precuneus compared to control (**Supplemental Figure 5B**, *p* < 0.01) and asymptomatic (**Supplemental Figure 5B**, *p* < 0.05) cases. There was an elevation of Msn levels in AsymAD precuneus compared to controls, although it was not statistically significant. Lastly, since Msn expression is observed in microglia proximate to Aβ plaques in human AD and 5xFAD mouse brains, we sought to determine Msn protein abundance in a published proteomic analysis of laser-micro-dissected Aβ plaques from the hippocampus and adjacent entorhinal cortex of rapidly progressive AD (rpAD) and sporadic AD (spAD) cases [17]. We found Msn abundance to be significantly lower in rpAD cases compared to spAD cases (**Supplemental Figure 5C**, *p* < 0.001) showing that despite the observed increased Msn expression in AD, it appears to have a negative association with rapidly progressive AD.

**Figure 5.**
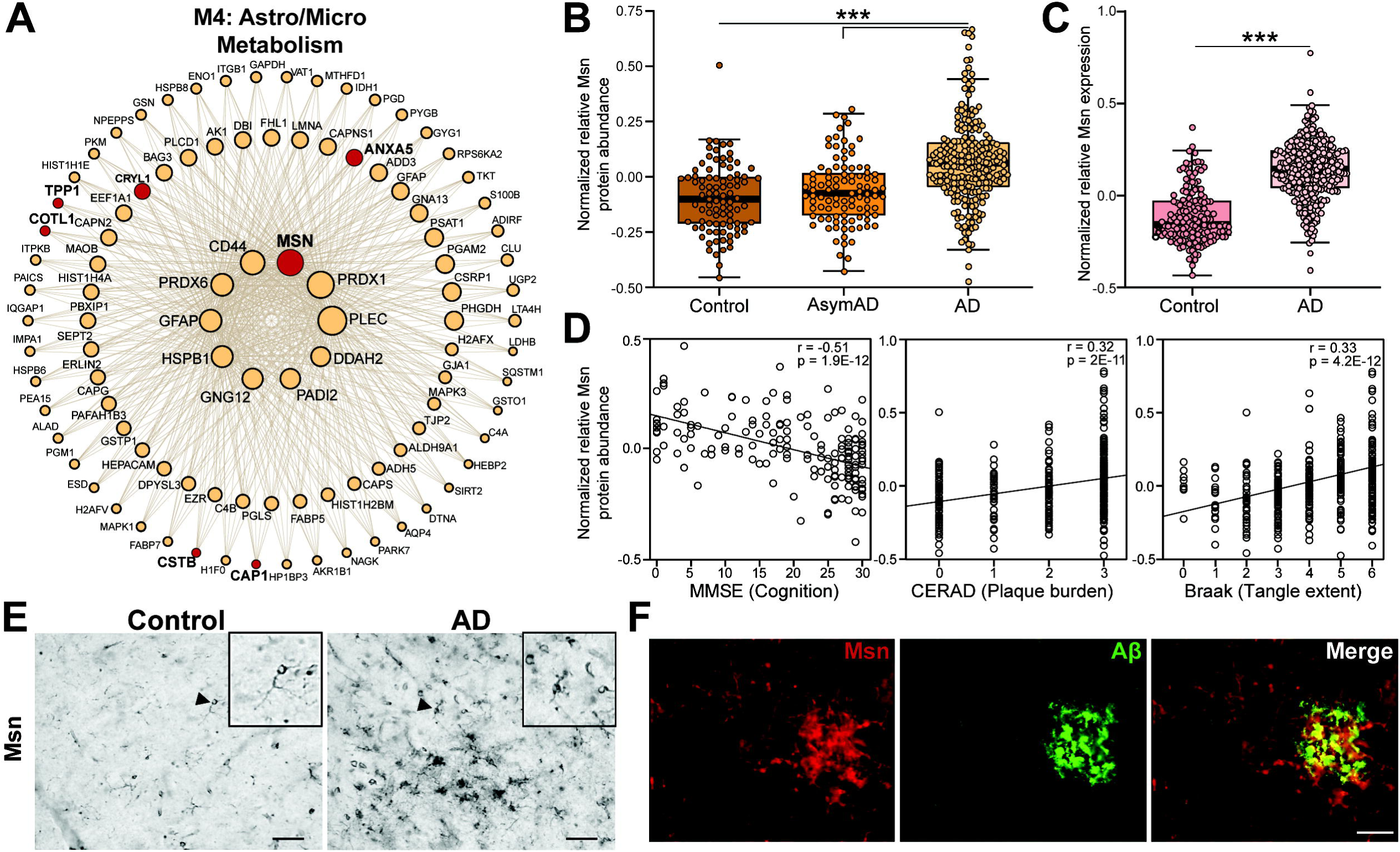
Moesin is a hub protein of an AD-associated proteomic module with increased expression in the human AD brain. **A** Msn was identified as a hub of a microglial-enriched protein module identified by weighted co-expression network analysis (WGCNA) of proteomic data from controls, asymptomatic AD, and AD human post-mortem brain (dorsolateral prefrontal cortex) [40]. The top 100 proteins by module eigenprotein correlation value (kME) in the M4 module are shown. The size of each circle indicates the relative kME. Red circles indicate known microglia-enriched proteins identified in the current proteomic analysis. **B** Normalized relative Msn protein abundance in controls (*N* = 91), AsymAD (*N* = 98), and AD cases (*N* = 231) human post-mortem brains. Protein abundance data obtained from a published proteomic dataset [40]. One-way ANOVA, Tukey *post hoc* analysis: \*\*\**p* <□ 0.001. **C.** Normalized Msn mRNA expression in controls (*N* =157) and AD (*N* = 308) human post-mortem brains (dorsolateral prefrontal cortex). RNA expression data obtained from a published dataset [42]. Unpaired t-test: \*\*\**p* <□0.001. **D** Scatterplots displaying correlation between Msn protein abundance in human post-mortem brain and cognitive function (MMSE, mini-mental status examination score, higher scores represent better cognitive function) of control (*N* = 26), AsymAD (*N* = 28), and AD (*N* = 83) cases, amyloid-beta (Aβ) plaque burden (CERAD, Consortium to Establish a Registry for Alzheimer’s disease Aβ plaque score, higher scores represent greater plaque burden) in control (*N* = 91), AsymAD (*N* = 98), and AD (*N* = 230) cases, and tau tangle extent (Braak, tau neurofibrillary tangle staging score, higher scores represent greater extent of tangle burden) in control (*N* = 91), AsymAD (*N* = 98), and AD (*N* = 230) cases. Correlation coefficient (R) and significance of the association are shown. **E** Representative immunohistochemical images of control (*N* = 10) and AD (*N* = 10) post-mortem human brains (frontal cortex) stained with Msn. The inset, indicated by arrowhead, displays a high magnification of the Msn positive cells with ramified microglial morphology in control brain and activated microglial morphology in AD brain. Scale bar = 40µm. **F** Representative immunofluorescence images of human AD brain (frontal cortex) showing Msn (red) and Aβ (green). Scale bar = 50µm.

In addition to exploring human proteomic datasets, we examined published TMT-MS and RNAseq data for Msn protein and transcript levels, respectively, in mouse models of AD pathology [43, 44]. Quantitative proteomics using TMT-MS was conducted on brains from WT, 5xFAD (Aβ pathology), JNPL3 (tau pathology), and a cross of 5xFAD and JNPL3 (Aβ & tau pathologies) mice at 4 months, 7 months, and 10 months of age. There was an age dependent increase in Msn protein abundance in the 5xFAD and 5xFAD/JNPL3 brains compared to age-matched WT brains (**Supplemental Figure 6A**, *p* < 0.001), while Msn protein abundance was unchanged in JNPL3 mice. We observed a similar age dependent increase in Msn transcript levels in 5xFAD brain compared to WT brains, with a significant increase at 12 months of age (**Supplemental Figure 6B**, *p* < 0.05). Overall, analysis of these independent human and mouse proteomic and transcriptomic studies increases the validity of our finding of increased Msn in human AD and highlights a potential novel role for Msn in FTLD-TDP pathology.

## Discussion

Studies of human brain gene and protein expression have consistently identified microglial genes/proteins within immune pathways as determinants of disease progression and cognitive decline [3, 30, 31]. Molecular characterization of microglia has been traditionally biased towards transcriptomic studies [42, 57, 58] rather than proteomic studies due to the generally low protein yield from isolated cells, challenges related to microglial isolation from the brain, and technical requirements for mass spectrometry analyses. Multiple studies have shown the discordance between transcript-level and protein-level expression data attributed to post-transcriptional processes such as post-transcriptional mRNA regulation, post-translational protein modifications, protein recycling and degradation [19–21]. Therefore, comprehensive profiling of proteins, rather than transcripts, of microglial cells is necessary for a deeper understanding of microglia-mediated disease mechanisms in neurodegenerative diseases such as AD. Prior proteomic studies have used magnetic-activated cell sorting (MACS) enrichment strategies to isolate microglia from fresh mouse or human brain [22, 25, 59]. MACS-enrichment aims to facilitate a rapid, high-throughput, immuno-magnetic separation of a pure population of a desired cell-type, i.e. microglia, from a single cell-suspension; however, it is limited by contamination by non-microglial proteins despite high cellular purity.

In this first FACS-based microglial proteomics study in adult wild-type mice, we have established the feasibility of this approach for microglial proteomics and have demonstrated that FACS-based microglial enrichment is the approach of choice for microglial proteomics studies. As compared to the MACS strategy, FACS effectively gates out protein-rich debris in the preparation which are likely cellular remnants of other cells such as neurons, astrocytes and oligodendrocytes that are enriched along with microglia due to the cellular complexity of the brain. Since non-cellular elements are likely to be non-nucleated, FACS or MACS approaches can be equivalent approaches for transcriptomic studies due to high cellular purity. However, proteomics studies are easily skewed by proteins in non-cellular elements in the preparation. As a result of this advantage of FACS, the proteomes obtained by each isolation strategy are indeed very different: the MACS-enriched microglia proteome over-represents synaptic proteins, suggesting a significant neuronal component in these samples, while the FACS-isolated microglia proteome is enriched for immune function proteins, indicating higher enrichment of microglial proteins. In order to assess whether the FACS proteome was indeed enriched for microglial proteins, we performed cell-type enrichment analysis of the differentially expressed proteins between the MACS and FACS proteomes and demonstrated that the FACS-isolation approach, when coupled with TMT-MS, is a superior method yielding a proteome that is highly enriched for canonical microglia-specific proteins while non-microglial proteins, particularly those derived from neurons and oligodendrocytes, are significantly depleted.

A strength of our study is the use of TMT-MS for quantitative proteomic characterization. The multiplex paradigm enables the accurate quantitation of thousands of proteins across many samples simultaneously for large-scale quantitative proteomic applications [60, 61]. Also, a major advantage of TMT-MS is the ability to multiplex all of the peptide sets prepared from multiple samples to be combined into a single LC-MS/MS analysis, resulting in improved breadth of coverage by minimizing missing values that are common in ‘shotgun’ label-free based quantification [28, 62]. The combination of TMT-MS and SPS-MS3 methods significantly improved the acquisition, and quantitation of MACS-enriched and FACS-isolated microglial proteomes. Critically, collapsing the 10-TMT channels (5 samples/isolation strategy) resulted in higher signals and identified proteins, overcoming the challenges of low protein yield typically expected from isolation of rare cell-types such as microglia. Many studies pool isolated microglia from 3-12 mouse brains for a MS run to increase protein yield and improve detection/quantitation of proteins in a MS run [11, 22, 25]; however, each sample in our study consisted of microglial cells isolated from one whole mouse brain without pooling for either isolation strategy. This is significant because 1) we demonstrate that at least 18,000 FACS-isolated microglia from one mouse brain is sufficient for a thorough proteomic analysis, 2) sampling bias is minimized when samples are not pooled, and 3) this method is especially cost-effective for complex, long-term mouse studies. The limitations of our approach are also made apparent by the aforementioned points. We identified approximately 1,800 proteins compared to the 4,133 proteins identified in our previous proteomic analysis of acutely isolated microglia [25]. This low number of identified proteins could have been ameliorated by pooling brain samples, as this significantly increases net protein yield. Additionally, enzymatic digestion rather than mechanical dissociation of brain tissue prior to isolation could increase microglial cell yield from one whole brain, and thus, protein yield. Notably, mechanical subcellular dissociation coupled to density-based centrifugation for purification of intact nuclei from brain for various downstream applications including FACS-MS has been demonstrated to achieve high purity separation of neuronal from non-neuronal nuclei [63] and this approach could incorporate detection of nuclear microglial-specific marker proteins from the current study to purify microglial nuclei. Lastly, the increased number of protein isoforms found by MACS could be attributed to non-cellular contamination in the MACS-only samples.

Our study also supports the use of our validated pipeline of FACS-based isolation coupled with TMT-MS for characterization of non-microglial brain cell types. One advantage of the FACS approach is that it provides the opportunity to isolate other cell-types from the brain, which is not feasible using MACS approaches without compromising cell integrity and viability. In our current study, we did not perform enzymatic digestion of the brain to maximize microglial enrichment prior to FACS or MACS. Enzymatic digestion can significantly increase the yield of endothelial cells as well as other glial cells (astrocytes and oligodendrocytes), allowing simultaneous isolation of multiple cell types by FACS. Recently, a cell isolation methodology termed concurrent brain cell type acquisition (CoBrA) has been used to isolate microglia, endothelial cells, astrocytes, and oligodendrocytes from mouse brain for RNAseq studies [64]. Unlike transcriptomic approaches, proteomic applications require specific strategies to reduce protein contamination from serum, albumin, and other proteins such as keratin, which we are currently optimizing for an analogous simultaneous cell-type isolation pipeline for proteomic studies. Based on our demonstration of a high-quality proteome from FACS-isolated microglia, which traditionally have very low protein yield per cell, our results support the feasibility of using FACS to isolate distinct cell types with high purity while minimizing non-specific contamination, for simultaneous proteomic characterization of multiple cells types. This is particularly important for resolving why we observed a significant enrichment of endothelial genes/proteins in our FACS-isolated microglial proteome, which could mainly be a reflection of our cell-type enrichment analysis method for endothelial cells. The proteomic reference markers suffer from the absence of an endothelial proteomic profile, by which markers that are common to both microglia and endothelia could be excluded by the thresholding rubric performed to define the cell-type enriched marker lists. Current brain endothelial cell biology is solely based on expression at the transcript level because no purified endothelial proteomes exist, thus, our cell-type enrichment analysis was conducted using a mouse brain cell-type transcriptomic reference [11]. Given the discordance between transcript-level and protein-level expression, it is difficult to conclude that there is indeed endothelial cell-type protein enrichment by FACS targeting microglial enrichment. This can only be resolved by proteomic characterization of concurrently isolated microglial cells and endothelial cells from the same mouse brain and comparison of these data with existing endothelial transcriptomic data.

We assessed our MACS-and FACS-microglial proteomes for cell-type enrichment with two different mouse brain cell-type reference datasets: cell-type resolved proteome by Sharma *et al.* [22] and cell-type resolved transcriptome by Zhang *et al*. [11] Although, both reference datasets identified the FACS-proteome to be enriched with microglial proteins/gene symbols and depleted of non-microglial proteins/gene symbols, there were still a significant number of gene products that were not assigned to a specific cell-type. This could be attributed to several factors. First, it could be that these proteins/genes are ubiquitous in the brain regardless of the cell-type. Second, there are inherent differences in the isolation methodologies between our study and the reference studies. For example, the reference cell-type proteome [22] isolated cells using MACS-enrichment, while the reference cell-type transcriptome [11] utilized FACS isolation from transgenic mice and immunopanning to isolate cells. Lastly, the expression or regulation of these proteins/gene symbols in their respective cell-type might vary with age. We characterized the microglial proteome of adult mice, while both reference datasets characterized microglia and other brain cell-types from vastly younger mice ranging in age from P1 to P8* [11, 22]. In summary, the combination of all these factors may have limited our ability to fully resolve the cell-type origin of the proteins identified within our dataset and highlights the need for comprehensive characterization of the proteome or transcriptome of mouse brain cell-types using consistent mouse models at similar ages, isolation protocols, and technical and analytical measures that will allow for cross-study comparisons.

Despite the advantages of the FACS approach, highly-abundant microglia proteins should still be identified using either strategy. Therefore, using a consensus between both approaches, we identified several proteins with high abundance in both datasets, and the top 10 proteins of this list were expressed in both datasets at or above the 90^th^ percentile of abundance. Of these highly-abundant and microglia-specific proteins, we considered Msn and Cotl1 for validation studies. Cotl1 protein expression was observed in microglia of WT and 5xFAD brains, with higher immunofluorescence and increased microglial size observed in the 5xFAD mice only, consistent with our prior observations [25]. The findings pertaining to Msn represent novel observations for this protein in mouse models of AD pathology and human AD. In WT mice, Msn was expressed in ramified/homeostatic microglia as well as non-microglial cells that resembled endothelial cells, consistent with prior transcriptomic studies showing that Msn is also highly abundant in endothelial cells [11]. In the 5xFAD mouse brains, Msn protein expression was localized to large clusters of reactive microglia rather than ramified microglia or endothelial cells, specifically within microglial processes, that surround and infiltrate Aβ plaques, suggestive of the disease-associated microglia (DAM) phenotype that was identified previously [13]. Similar to our findings in 5xFAD brains, we also found increased Msn expression in the vicinity of Aβ plaques in human AD brain. Consistent with ezrin-radixin-moesin (ERM) proteins, the pattern of Msn expression in microglia appeared to be more membrane localized rather than cytosolic [65]. These immunohistochemical studies show that Msn and Cotl1 proteins, like other known markers such as Tmem119 or Iba1, could serve as novel markers of microglia in the mouse brain in normal and disease states.

In a large human brain network built using over 400 frontal cortex samples, Msn was found to be a hub of a protein co-expression module that was strongly associated with neuropathology and cognitive decline, indicating a key role for Msn in AD pathogenesis. Furthermore, we found Msn protein abundance was significantly elevated in AD DLPFC and precuneus. We also found that Msn around plaques is lower in rapidly progressive AD brains as compared to sporadic AD brains. Our *in-vitro* data, combined with correlative data from human brain studies, support the idea that Msn in microglia may play important roles in AD pathogenesis by impacting phagocytosis and pro-inflammatory cytokine release. We identified that siRNA-mediated knockdown of Msn augmented the release of TNF and IL-10 by primary mouse microglia in response to LPS stimulation TNF or tumor necrosis factor is a pro-inflammatory molecule [66, 67] while IL-10, although traditionally considered anti-inflammatory, has been found to have detrimental effects in AD models [68, 69]. While we observed an increase in inflammatory cytokine release by microglia following Msn knockdown, we also observed an increase of Msn in AD cases, and increase in TNF in AD has been previously demonstrated [68]. One very likely explanation for this apparent discrepancy between our *in-vitro* studies and human AD findings is that we used a pure microglial culture system without any other cell-types that might impact TNF levels or be a source of TNF itself. Indeed, in the brain, there are multiple sources of TNF, such as astrocytes, sub-populations of neurons, and even different populations of microglia [69–72]. Furthermore, these experiments do not account for the microglial heterogeneity observed in *in-vivo* AD models [73], making it difficult to know whether loss of Msn in this context will also impact TNF release in a similar manner. Msn silencing also decreased the ability of microglia to phagocytose Aβ fibrils. The role of Msn in phagocytosis is not unexpected because Msn belongs to the family of ERM proteins which are involved in actin-associated biological processes [49, 51, 52, 65]. Altogether, these findings raise the possibility that increased expression of Msn in AD may represent a compensatory and potentially protective mechanism that is most likely explained by increased microglial activation. Future studies using conditional Msn deletion in microglia or endothelial cells in mouse models of AD pathology will clarify the protective or detrimental role of Msn in AD pathogenesis. Based on our *in-vitro* findings of Msn deletion in microglia, additional mechanistic studies are also needed to define the molecular and cellular defects responsible for impaired Aβ phagocytosis in Msn-deficient microglia.”

## Conclusion

In conclusion, this study establishes the feasibility and superiority of the FACS-based TMT-MS approach for quantitative proteomics of acutely-isolated microglia from adult mouse brain which can now be applied for future quantitative proteomic studies of adult mouse microglia using disease models. Using this strategy, we defined a core set of highly abundant microglia-specific proteins. Among these, we provide novel evidence for a significant and potentially protective role for Msn in human AD.

## Abbreviations

Aβ: amyloid-beta
ACN: acetonitrile
AD: Alzheimer’s disease
ANOVA: analysis of variance
AGC: automatic gain control
AsymAD: asymptomatic Alzheimer’s disease
BCA: bicinchoninic acid
CE: collision energy
COBRA: concurrent brain cell type acquisition
DAM: disease-associated-microglia
DAPI: diamidino-2-phenylindole
DTT: dithiothreitol
ER: endoplasmic reticulum
ERM: ezrin-radixin-moesin
FDR: false discovery rate
FBS: fetal bovine serum
FACS: fluorescence activated cell sorting
FA: formic acid
fAβ42-488: fibrillar fluorescent Aβ42 conjugated to HiLyte Fluor 488
GO: gene ontology
GWAS: genome-wide association studies
HCD: higher energy collision dissociation
IAA: iodoacetamide
KD: knockdown
LFQ: label-free quantitation
LPS: lipopolysaccharide
MACS: magnetic-activated cell sorting
MS: mass spectrometry
MMSE: mini-mental status examination
Msn: moesin
PBS: phosphate buffered saline
PE: phycoerythrin
PSM: peptide spectral match
qRT-PCR: quantitative reverse transcriptase pcr
rpAD: rapidly progressive ad
RT: room temperature
SNPs: single-nucleotide polymorphisms
siRNA: small interfering ribonucleic acid
sAD: sporadic ad
SIP: stock isotonic percoll
SPS: synchronous precursor selection
TMT: tandem mass tag
TEAB: triethylammonium bicarbonate
TFA: triflouroacetic acid
TBS: tris buffered saline
WGCNA: weighted correlation network analysis

## Declarations

### Ethics approval and consent to participate

Approval from the Emory University Institutional Animal Care and Use Committee was obtained prior to all animal-related studies (IACUC protocol # PROTO201800252).

### Consent for publication

All authors have approved of the contents of this manuscript and provided consent for publication.

### Availability of data and materials

The mass spectrometry proteomics data have been deposited to the ProteomeXchange Consortium via the PRIDE partner repository with the dataset identifier PXD015652.

### Authors’ contributions

#### Conceptualization

NTS, S.Rangaraju

#### Methodology

S.Rayaprolu, TG, HX, S.Ramesha, LDW, JS, DMD, EBD, JAW, RB

#### Investigation

S.Rayaprolu, NTS, S.Rangaraju

#### Writing-Original draft

S.Rayaprolu, NTS, S.Rangaraju

#### Writing-Review and Editing

TG, HX, S.Ramesha, LDW,JS, DMD, EBD, JAW, JJL, LBW, RB, AIL

#### Funding acquisition

S.Rayaprolu, LBW, AIL, NTS, S.Rangaraju

#### Resources

LBW, JJL, AIL, NTS, S.Rangaraju

#### Supervision

NTS, S.Rangaraju

All authors read and approved the final manuscript.

## Acknowledgements

Research reported in this publication was also supported in part by the Emory Flow Cytometry Core (EFCC) and Emory University Integrated Cellular Imaging (ICI) Microscopy Core, two of the Emory Integrated Core Facilities (EICF). EFCC is subsidized by the Emory University School of Medicine and additional support was provided by the National Center for Georgia Clinical & Translational Science Alliance of the National Institutes of Health under Award Number UL1TR002378. ICI Microscopy core is subsidized by the Emory University Integrated Cellular Imaging Microscopy Core of the Emory Neuroscience NINDS Core Facilities grant, 5P30NS055077. The content is solely the responsibility of the authors and does not necessarily reflect the official views of the National Institute of Health.

## Additional Files

**Table S1.** Quantitative protein expression data from MACS-enriched and FACS-isolated mouse microglia. Log_2_ transformed protein abundance data and differential expression analysis.

**Table S2.** GO Elite analysis of 953 significantly differentially expressed proteins between MACS-enriched and FACS-isolated mouse microglia proteomes.

**Supplemental Figure 1. Viability of mechanically dissociated mouse brain mononuclear cells.** Representative flow cytometry data displaying isolation of >95% live CD11b^+^ microglia from mechanically dissociated fresh, whole mouse brian (*N* = 4) following percoll density centrifugation.

**Supplemental Figure 2. Enrichment of microglial and endothelial specific proteins by FACS. A-D** Histograms displaying top 10 differentially expressed **A** microglia, **B** neuron, **C** astrocyte, and **D** oligodendrocyte cell-type proteins (defined by Sharma *et al.* [22]) in FACS-isolated microglia proteome and MACS-enriched microglia proteome. The Y-axis shows list of proteins, X-axis shows Log_2_-transformed normalized abundance (abundance/row geomean), and error bars represent ± SEM. **E** Volcano plot displaying the distribution of differentially expressed proteins between FACS-isolated and MACS-enriched microglia proteomes. Cell-type enrichment defined by a reference cell-type transcriptome, Zhang *et al.* [11], shows significant enrichment of microglial and endothelial specific proteins in the FACS proteome (*p* < 0.05, Unpaired t-test). Red dots = microglia, turquoise dots = neuron, pink dots = astrocyte, yellow dots = oligodendrocyte, green dots = endothelial cell. Grey dots represent differentially expressed proteins with a *p* > 0.05. Log_2_ fold-change is shown on the X-axis, -Log_10_(*p*-value) is shown on the Y-axis, and horizontal dotted line indicates *p* = 0.05. **F** Venn diagrams comparing number of cell-type specific proteins defined by our two cell-type enrichment analyses: reference cell-type proteome: Sharma *et al.* [22] and reference cell-type transcriptome: Zhang *et al.* [11].

**Supplemental Figure 3. Contamination of non-microglial proteins in MACS-enriched microglia proteome**. **A** Pie chart displaying distribution of cell-type enrichment in previously published microglial proteome, Rangaraju *et al.* [25], where 4.5% of the 4,133 quantified proteins are microglial-specific while nearly 17% of the proteins are from other brain cell-types. **B** Highly-abundant proteins in a MACS microglial proteomic study [25] included non-microglial proteins such as Mbp, Aldoa, Gfap, and Camk2a. Micro = microglia, Neu = neuron, Astro = astrocyte, Oligo = oligodendrocyte.

**Supplemental Figure 4. Validation of Cotl1 in microglia. A** Representative immunofluorescence images of Cx3cr1^CreER-YFP^-WT (*N* = 4) and Cx3cr1^CreER-YFP^-5xFAD (*N* = 6) mouse cortex stained for GFP (microglia) and Cotl1. Arrow indicates microglia immunopositive for GFP (to detect microglia) and Cotl1. **B** Representative images of 9-10 month old WT and 5xFAD cortex used for quantitative analysis of Msn fluorescence intensity shown in a histogram below. Unpaired t-test, \*\*\*\**p* < 0.0001.

**Supplemental Figure 5. Moesin protein levels are increased in human AD and FTLD-TDP brain. A** Msn protein abundance in dorsolateral prefrontal cortex post-mortem brain tissue of control (*N* = 43), AD (*N* = 47), FTLD-TDP (*N* = 29), ALS (*N* = 54), and PD/PDD (*N* = 76) cases; UPENN cohort. Protein abundance was measured with LFQ [40]. One-way ANOVA, Tukey *post hoc*: \**p* <□0.05,\*\*\**p* <□0.001. **B** Msn protein abundance in precuneus post-mortem brain tissue of control (*N* = 13), AsymAD (*N* = 13), and AD (*N* = 20) cases; BLSA cohort. Protein abundance was measured with LFQ [40]. One-way ANOVA, Tukey *post hoc*: \**p* <□0.05,\*\*\**p* <□0.001. **C** Msn protein abundance in laser capture microdissected Aβ plaques from human post-mortem brain tissue of rapid progression AD (*N* = 22) and sporadic AD (*N* = 22) cases. Protein abundance was measured with LFQ by Drummond *et al.* [41]. Unpaired t-test: \*\*\**p* <□0.001.

**Supplemental Figure 6. Moesin protein and mRNA levels increase with age in 5xFAD brain. A** Normalized relative Msn protein abundance in brains of wild-type (WT), 5xFAD (Aβ pathology), JNPL3 (Tau pathology), and a cross of 5xFAD and JNPL3 (Aβ & tau pathologies) mice at 4 months, 7 months, and 10 months of age (*N* = 3/age/genotype). Protein abundance obtained by TMT-MS [43]. **B** Relative Msn expression, shown as FPKM values, in brains of WT and 5xFAD mice at 3 months, 6 months, and 12 months of age (*N* = 2-3/age/genotype) from a previously published study [44]. Error bars represent ± SEM. One-way ANOVA, Tukey *post hoc* analysis: \**p* <□ 0.05, \*\*\**p* <□ 0.001, \*\*\*\**p* <□ 0.0001.

