## Supplementary figures and images for "Flow-cytometric microglial sorting coupled with quantitative proteomics identifies moesin as a highly-abundant microglial protein with relevance to Alzheimer’s disease"

### Supplemental Figure 1

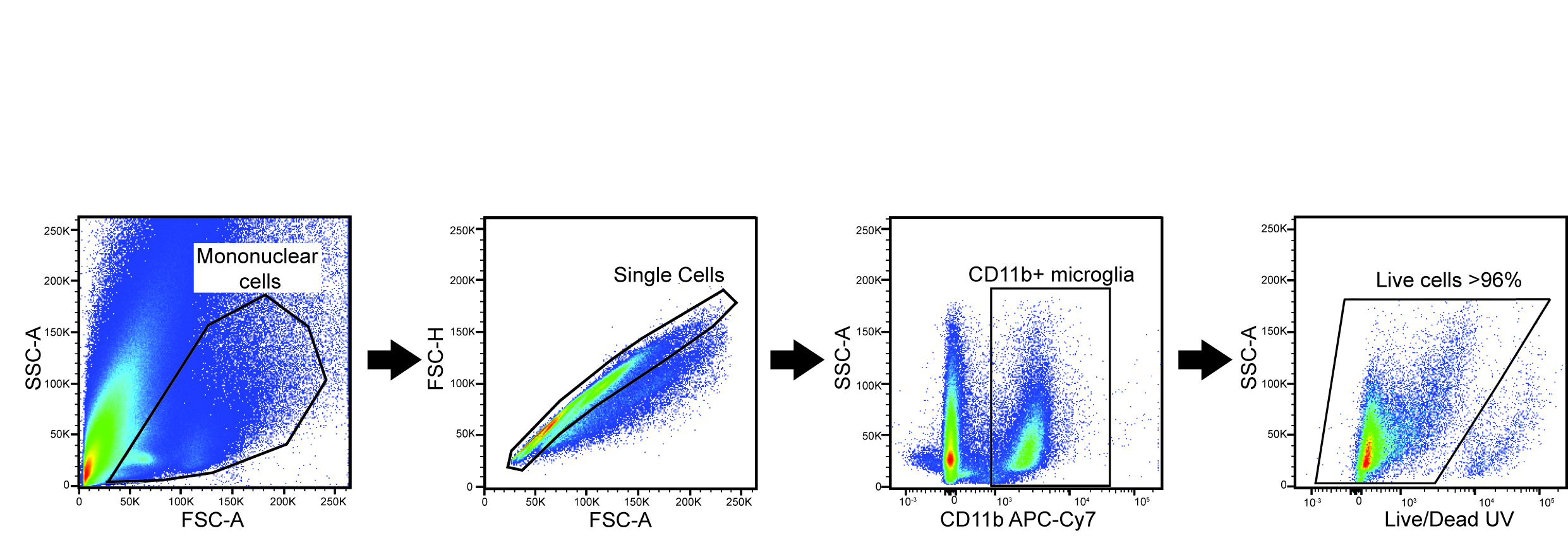

### Supplemental Figure 2

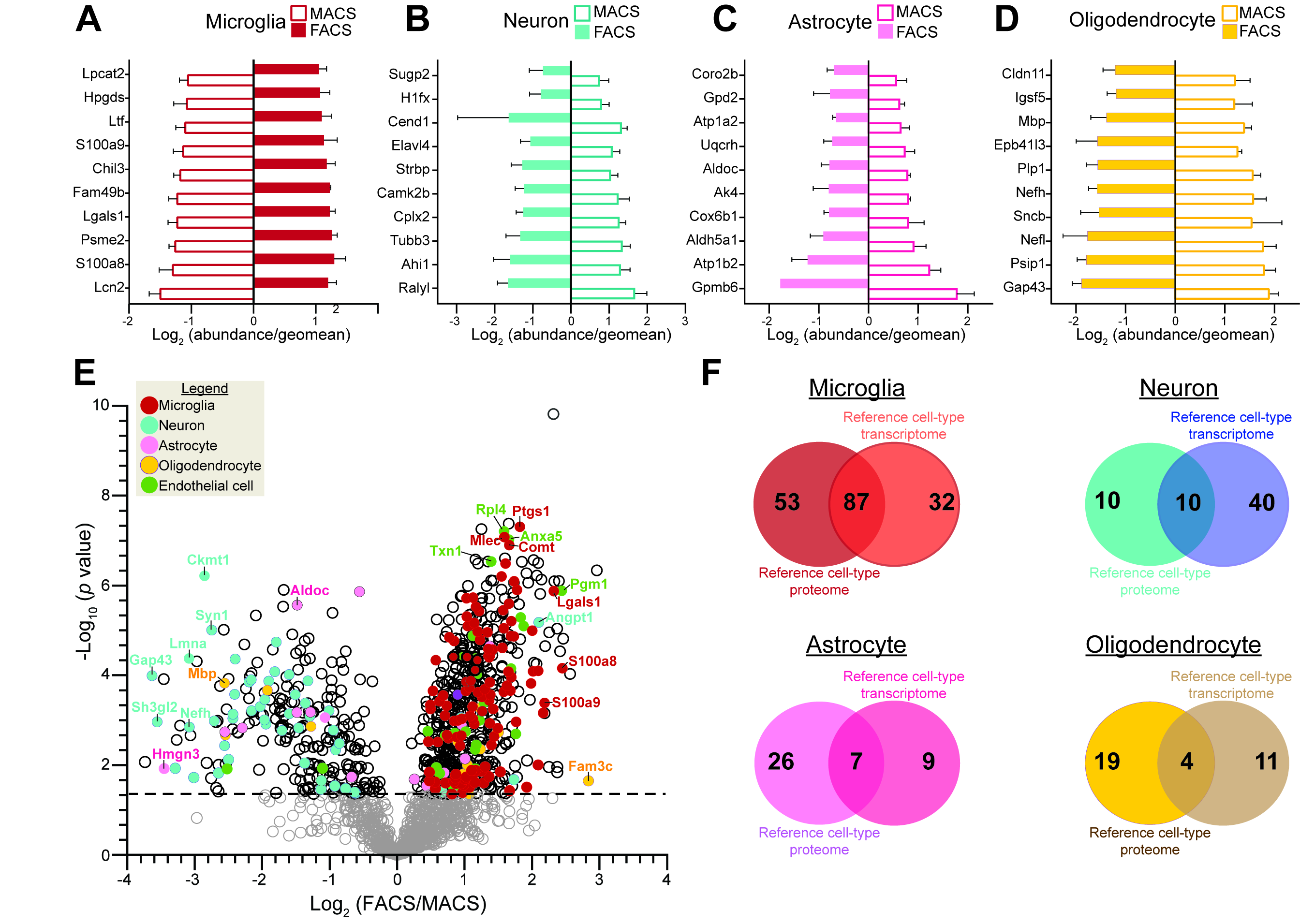

### Supplemental Figure 3

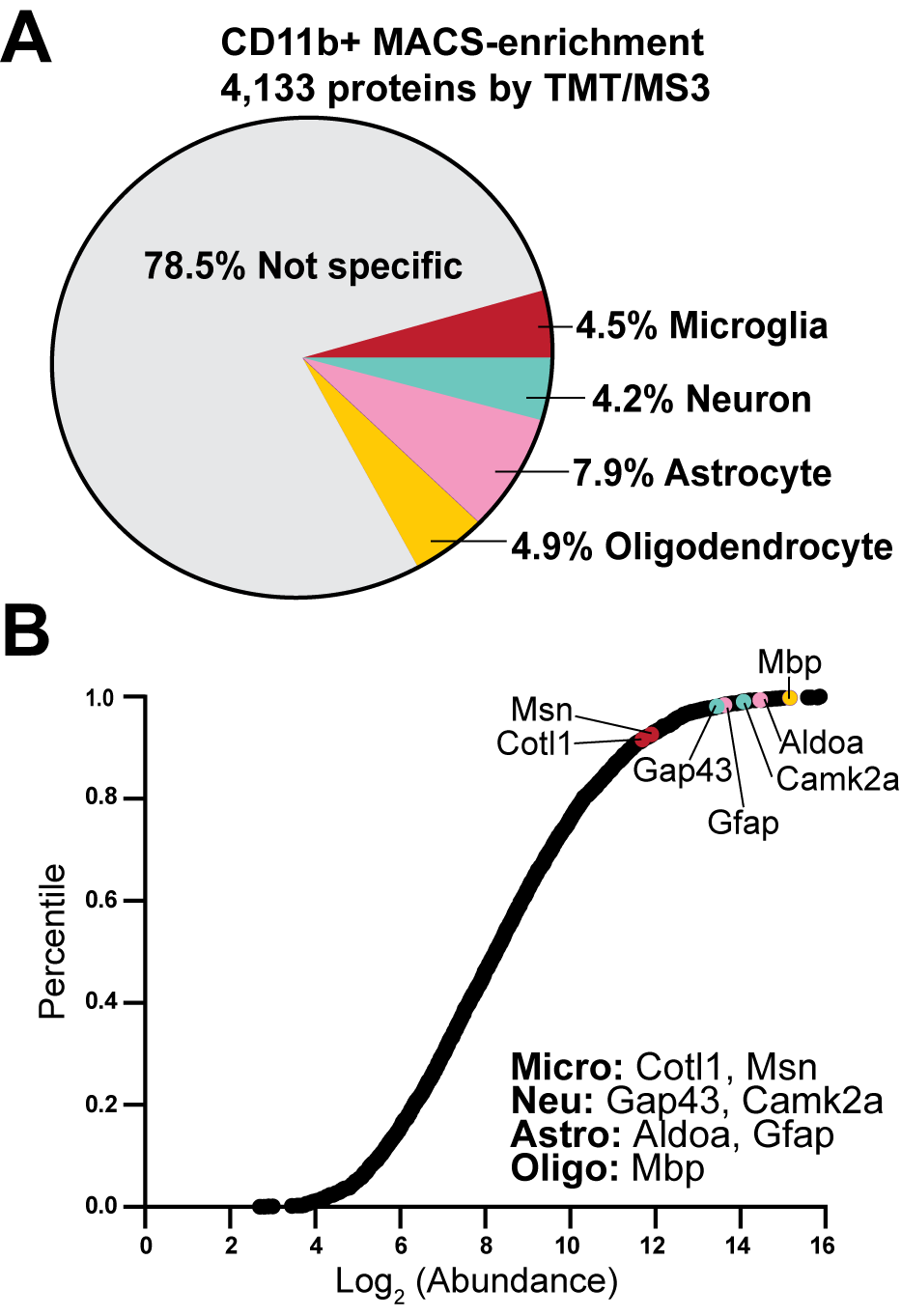

### Supplemental Figure 4

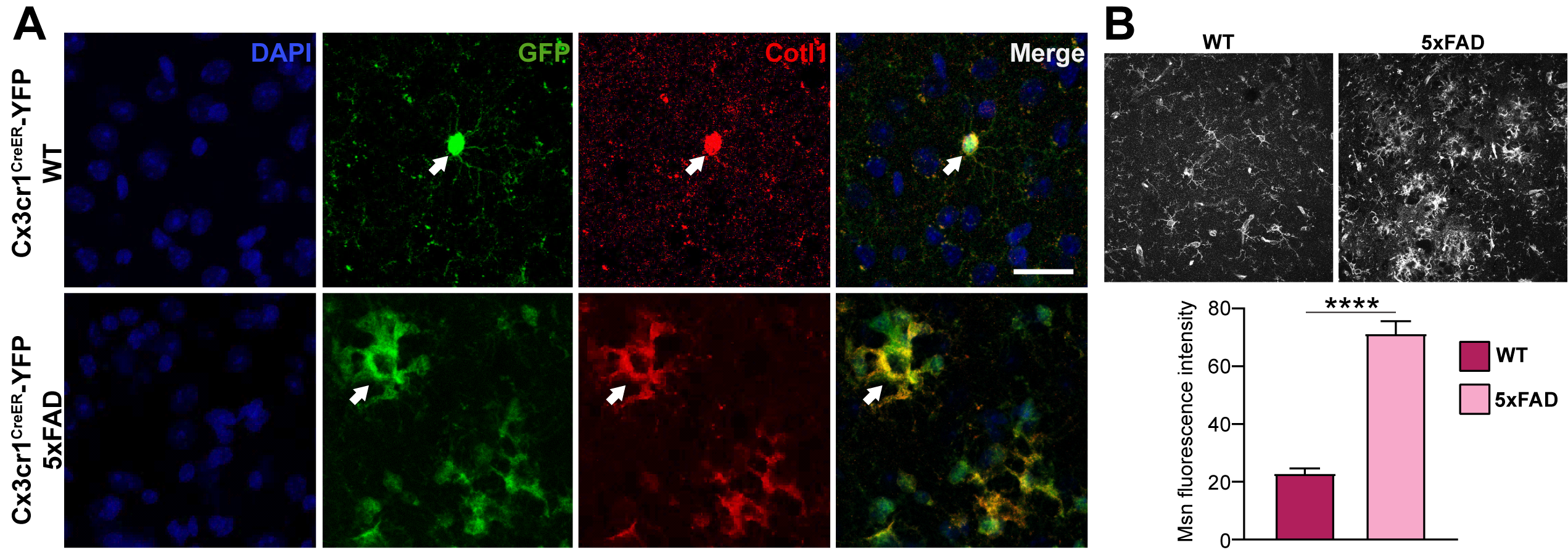

### Supplemental Figure 5

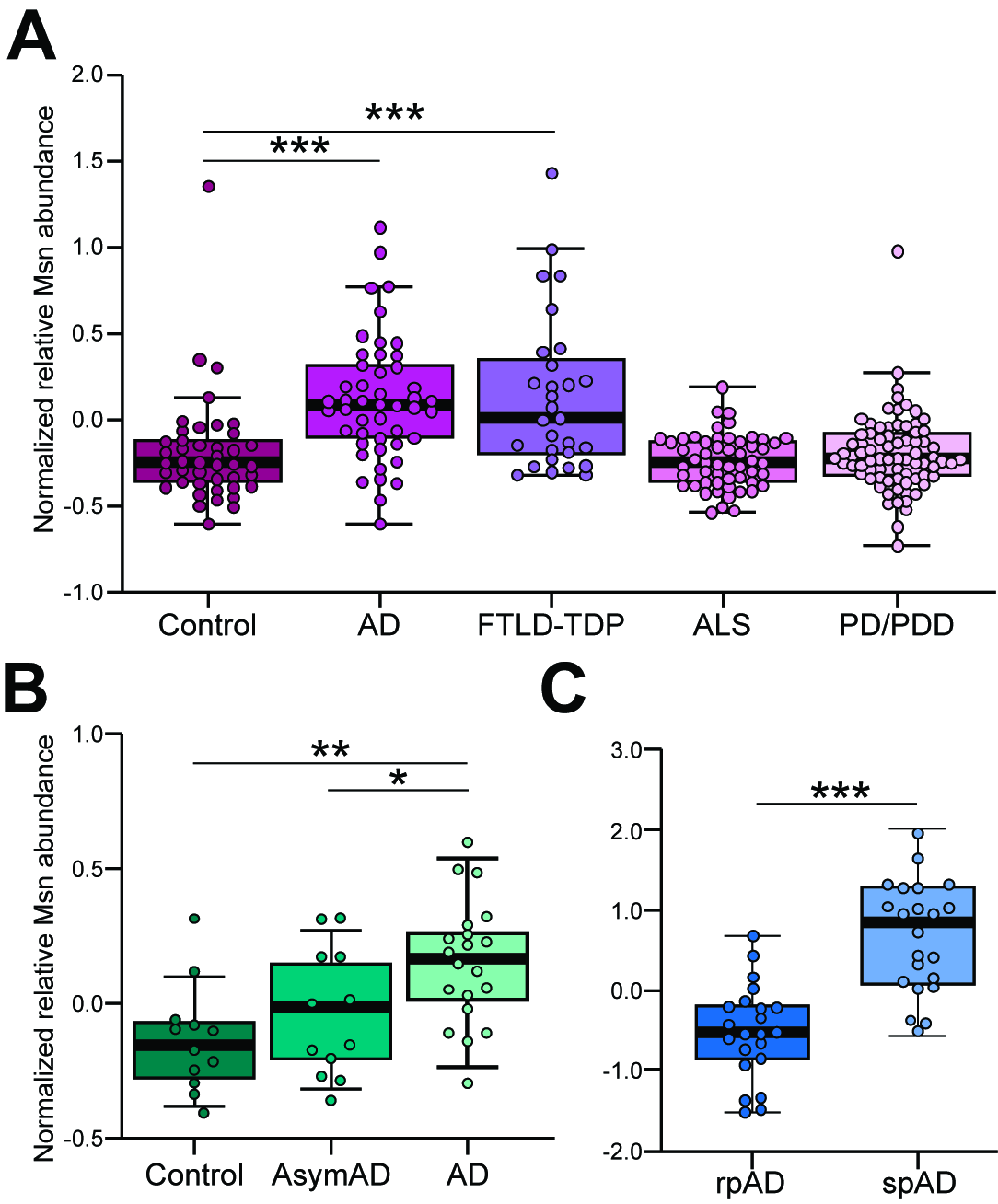

### Supplemental Figure 6

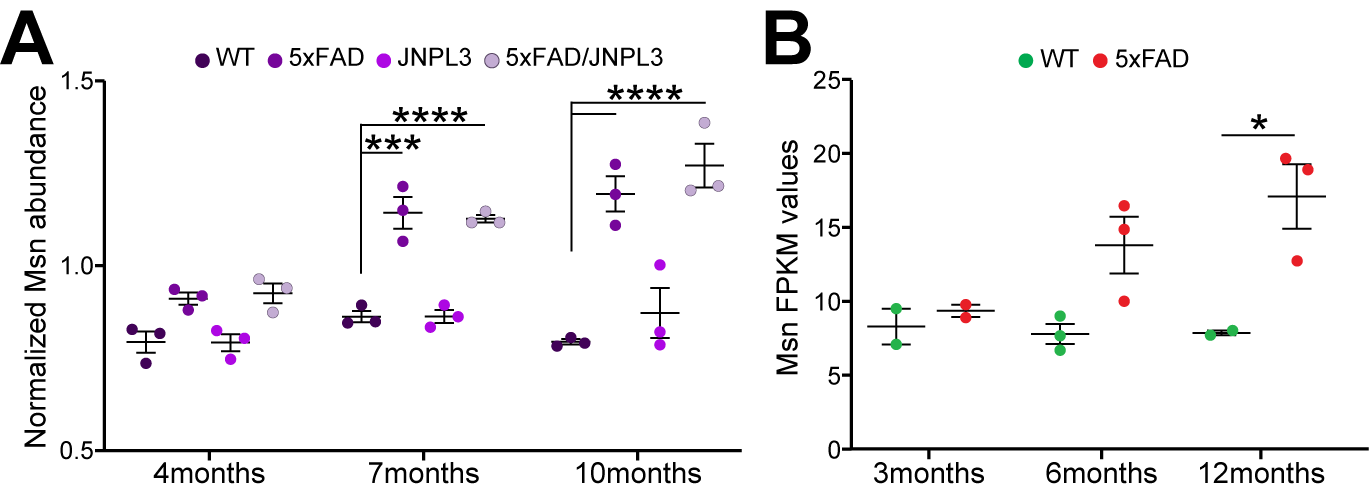
